# A lipid transport Mla Pqi Chimeric system is essential for *Brucella abortus* survival in macrophages

**DOI:** 10.1101/2024.10.31.621289

**Authors:** Adélie Lannoy, Alexi Ronneau, Miguel Fernández García, Marc Dieu, Patricia Renard, Antonia García Fernández, Raquel Condez-Alvarez, Xavier De Bolle

## Abstract

The envelope of diderm bacteria comprises of an inner membrane (IM) and an outer membrane (OM). Several pathways have been recently identified that facilitate the transport of phospholipids between the two membranes in *Escherichia coli*, including maintaining OM lipid asymmetry (Mla) and paraquat inducible (Pqi) systems. In this study, we report the identification and the characterization of a complex named Mpc in the intracellular pathogen *Brucella abortus*. Mpc is conserved in numerous species of Hyphomicrobiales and exhibits homology to both the Mla and Pqi systems. Mpc is essential for bacterial growth under conditions of envelope stress and for survival within macrophages during the early stages of infection. Analyses of protein-protein interactions and structural predictions indicate that the Mpc complex bridges IM to OM. The absence of this system results in an altered lipid composition of the OM vesicles, supporting the fact that Mpc plays a role in the transport of lipids between membranes. The discovery of a novel lipid trafficking system enhances the diversity and complexity of known lipid trafficking systems within diderm bacteria.

## Introduction

The integrity of the cell envelope is crucial for the survival and growth of bacteria. Gram-negative bacteria are characterised by the presence of an inner membrane (IM), a periplasmic space containing a thin layer of peptidoglycan (PG), and an outer membrane (OM). The IM is a bilayer of phospholipids (PLs) while the OM is an asymmetric bilayer. The OM inner leaflet is predominantly composed of PLs, while the outer leaflet lipid is mainly composed of lipopolysaccharide (LPS). The OM functions as an essential protective and permeability barrier (<u>Silhavy *et al*, 2010</u>).

All envelope components, whether proteic or lipidic, are biosynthesised in the cytoplasm or in the inner leaflet of the IM. They are then translocated from the cytoplasm to the outer leaflet of the IM, or flipped across the IM (<u>Silhavy *et al*., 2010</u>). In *Escherichia coli*, the transport systems across the periplasm are relatively well-understood with regard to the delivery of outer membrane proteins (OMPs) (<u>Konovalova *et al*, 2017</u>; <u>Tomasek & Kahne, 2021</u>), lipoproteins (<u>Grabowicz, 2018</u>) and LPS (<u>Giacometti *et al*, 2022</u>; <u>Okuda *et al*, 2016</u>) to the OM. However, the understanding of the systems which facilitate PLs transport are still unclear (<u>Kumar & Ruiz, 2023</u>).

Two distinct mechanisms are involved in the transport of PLs across the periplasm. Either proteins form a bridge spanning the entire periplasm, or a shuttle in the form of a soluble carrier protein is involved. The periplasmic-spanning AsmA-like protein family, which includes TamB, YhdP and YdbH, exhibits a specific conformation in a Taco-like shape. It is characterised by a hydrophobic groove, which facilitates the transport of lipids (<u>Cooper *et al*, 2023</u>; <u>Kumar & Ruiz, 2023</u>). Additionally, protein complexes spanning the periplasm with a channel shape have been described for Pqi (<u>p</u>ara<u>q</u>uat inducible) and Let (lipophilic <u>e</u>nvelope-spanning tunnel; formerly Yeb) (<u>Ekiert *et al*, 2017</u>; <u>Isom *et al*,</u> <u>2020</u>). The shuttle-based system is the Mla (<u>m</u>aintenance of OM lipid <u>a</u>symmetry) pathway (<u>Malinverni</u> <u>& Silhavy, 2009</u>). A common feature of the Mla, Pqi and Let systems is the presence of a protein comprising one or more MCE domains (<u>m</u>ediator of mammalian <u>c</u>ell <u>e</u>ntry), arranged as homo-hexamers in the structures that have been solved thus far (<u>Ekiert *et al*., 2017</u>; <u>Isom *et al*., 2020</u>; <u>Isom *et al*, 2017</u>; <u>Nakayama & Zhang-Akiyama, 2017</u>). The MCE superfamily of proteins is ubiquitous among diderm bacteria (<u>Arruda *et al*, 1993</u>; <u>Chen *et al*, 2023a</u>; <u>Cooper *et al*, 2024</u>; <u>Ekiert *et al*., 2017</u>; <u>Grasekamp *et al*, 2023</u>; <u>Isom *et al*., 2020</u>; <u>Isom *et al*., 2017</u>; <u>Malinverni & Silhavy, 2009</u>) and eukaryotic bacteria-derived organelles (<u>Awai *et al*, 2006</u>; <u>Isom *et al*., 2017</u>; <u>Lu & Benning, 2009</u>).

The Mla pathway is composed of the OmpC-MlaA complex in the OM, a MlaC periplasmic shuttle, that facilitates the transport of PLs through the periplasmic space, and MlaEFDB, a highly conserved <u>A</u>TP-<u>b</u>inding <u>c</u>assette (ABC) transporter complex in the IM. MlaD comprises a transmembrane helix and a periplasmic MCE domain and forms a ring-shaped homo-hexameric structure (<u>Coudray *et al*, 2020</u>; <u>Ekiert *et al*., 2017</u>). This system is responsible for the removal of mislocalised PLs from the OM and their subsequent transport back to the IM, which allows for the maintenance of the lipid asymmetry of the OM (<u>Chong *et al*, 2015</u>; <u>Malinverni & Silhavy, 2009</u>; <u>Shrivastava & Chng, 2019</u>). The Pqi complex is composed of an integral membrane protein of the IM (PqiA), devoid of an ATPase domain, and a PqiB homo-hexameric oligomer. This oligomer forms a stack of three rings composed of MCE domains which is followed by a syringe-like structure of coiled-coil alpha-helices (<u>Ekiert *et al*., 2017</u>). This structure spans the periplasmic space to interact with the lipoprotein PqiC in the OM. PqiC is arranged as a homo-octameric ring, which may serve to anchor the C-terminal part of PqiB to the OM, thereby stabilising the flexibility of the coiled-coil channel of the PqiB multimer (<u>Cooper *et al*., 2024</u>). The Let complex is composed of an integral membrane protein of the IM (LetA) lacking an ATPase domain, and LetB, a homo-hexameric oligomer containing a stack of seven MCE domains. The MCE domains form a channel that establishes an interaction between both membranes and capable of mediating direct lipid transport between the IM and OM (<u>Ekiert *et al*., 2017</u>; <u>Isom *et al*., 2020</u>).

It is noteworthy that MCE domains were initially identified in *Mycobacterium tuberculosis* (<u>Arruda</u> *<u>et al.</u>*<u>, 1993</u>).They were proposed to play a significant role in the transport of lipids and also in the virulence of bacterial pathogens (<u>Ekiert *et al*., 2017</u>). Furthermore, the integrity of the membranes is also a factor in the virulence of bacterial pathogens. The envelope structure and biogenesis of *Brucella abortus* were recently reviewed (<u>Alakavuklar *et al*, 2023</u>). *B. abortus* is a Gram-negative intracellular pathogen responsible for bovine brucellosis, a worldwide zoonosis with significant social and economic impacts (<u>Laine *et al*, 2023</u>; <u>Moreno & Moriyón, 2006</u>). In contrast to *E. coli*, the members of the Hyphomicrobiales order bacteria, such as *B. abortus*, *Agrobacterium tumefaciens* and *Sinorhizobium meliloti*, display a unipolar growth (<u>Brown *et al*, 2012</u>). The incorporation of LPS, PG and OMPs occurs at the new pole and the division site in *B. abortus* (<u>Servais *et al*, 2023</u>; <u>Vassen *et al*, 2019</u>). Moreover, the OM is linked to the PG by covalent cross-links between the N-terminus of OMPs and the peptide stems of PG (<u>Godessart *et al*, 2021</u>). Nevertheless, the biogenesis and turnover of the envelope in *B. abortus* remain under-investigated, particularly regarding the transport of PLs. As an intracellular pathogen, *B. abortus* encounters numerous stresses during its infectious cycle, such as antimicrobial peptides, which could potentially affect the integrity of its envelope.

In this study, we characterise the Mpc (Mla-Pqi chimeric) system comprising of the only MCE domain containing protein predicted from the *B. abortus* genome. The Mpc system is required for *B. abortus* growth in the presence of envelope stress and for survival during the first stage of macrophages infection. Structure prediction and pull-down analysis strongly suggest that the Mpc system forms a stable complex that spans the entire periplasmic space. The absence of the Mpc system generates an abnormal lipid composition of the OM vesicles, supporting its role in the trafficking of lipids between IM and OM in *B. abortus*.

## Results

### Identification of the *mpc* genes

In order to identify a system required to resist envelope stress, a transposon sequencing (Tn-seq) analysis was performed. A *Brucella abortus* 544 library (7.8 x 10^5^ random mutants), constructed with the mini-Tn*5* (<u>Sternon *et al*, 2018</u>), was grown on rich medium with 0.15 mg/mL of sodium deoxycholate (DOC) and compared to a control condition without DOC (Table S1). DOC, a detergent that affects the integrity of bacterial membranes (<u>Urdaneta & Casadesus, 2017</u>), was chosen over SDS and EDTA, which are commonly used in *E. coli*, because *Brucella spp*. are resistant to EDTA (<u>Moriyon & Berman, 1982</u>), but sensitive to DOC (<u>Caro-Hernandez *et al*, 2007</u>). To identify ORFs required for growth in the presence of DOC, the ΔTnIF variable was computed (Table 1). The more negative the ΔTnIF, the more the ORF is required for growth in the presence of DOC. The genes encoding for the BepCDE RND efflux pump, which is known to be involved in DOC resistance (<u>Martin *et al*, 2009</u>; <u>Posadas *et al*, 2007</u>) were selected here (Table 1), thereby confirming the efficiency of the Tn-seq approach for identifying genes involved in DOC sensitivity. The genes encoding for the CenKR two-component system (TCS), were identified with a ΔTnIF less than four times the genome standard deviation. This TCS involved in cell envelope integrity has been previously described in several Hyphomicrobiales (<u>Lakey *et al*, 2022</u>; <u>Skerker *et al*, 2005</u>) and *Brucella* spp (<u>Chen *et al*, 2023b</u>; <u>Liu *et al*, 2012</u>; <u>Zhang *et al*, 2009</u>). Of particular interest are the four more strongly genes (Table 1), with a ΔTnIF less than four times the genome standard deviation, which are found in the *mpcEFDA* operon. This operon encodes a potential system containing an MCE protein, which is involved in lipid transport (<u>Nakayama & Zhang-</u> <u>Akiyama, 2017</u>). This system has not yet been characterised in *Brucella* spp.

**Table 1.**
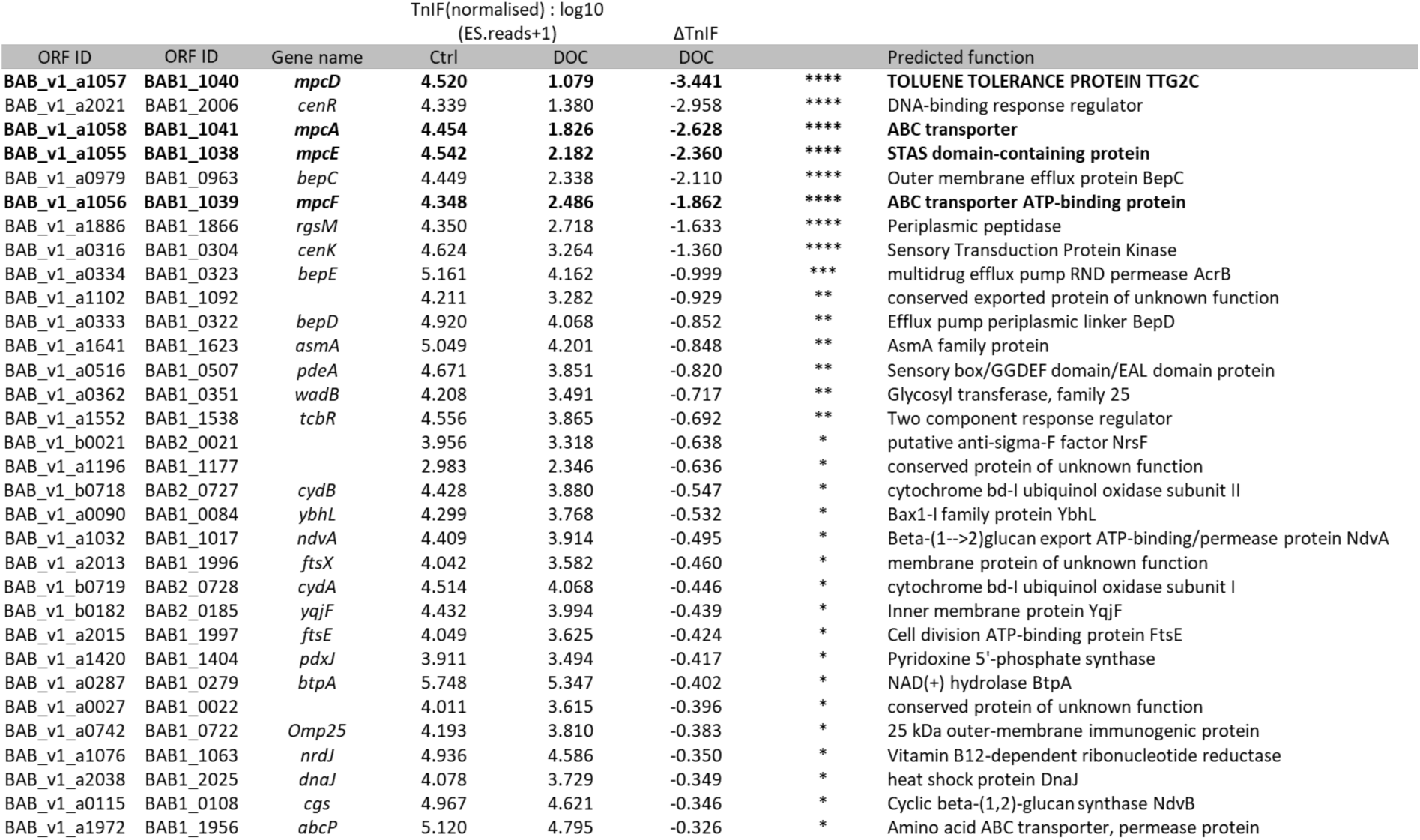
DOC-sensitive mutants according to Tn-seq. For each mutated gene, a Transposon insertion frequency (TnIF) is computed. Transposon insertion sites were identified through Illumina sequencing. In the control condition, there was an average of one unique insertion site every 2.62 bp, and in the envelope stress condition, every 2.48 bp, saturating the *B. abortus* genome in both conditions. Following reads mapping, we computed an ES.Reads value for each open reading frame (ORF), corresponding to the insertion number per bp for 80% of each ORF by excluding the first and last 10% of the predicted coding sequence. To enable a quantitative analysis of each ORF under different conditions, we calculated a transposon insertion frequency (TnIF) parameter. This frequency corresponds to the logarithm in base 10 of the mapped read numbers for the central 80% of each ORF (log_10_ (ES.Reads+1)). To identify genes required for growth in the presence of DOC (0.015%), we compared the TnIF of the stress condition to the control condition to obtain a ΔTnIF for each ORF (ΔTnIF = TnIF_DOC_ - TnIF_Ctrl_). The standard deviation (SD) of ΔTnIF values for the all the ORFs corresponds to 0.323. We considered ORFs with less than one SD (*; - 0.323) as less required, two SD (**; −0.647) as required, three SD (***; −0.97) as highly required and four SD (****; −1.293) as essential for growth on DOC.

To confirm the phenotype suggested by Tn-seq, markerless deletion strains for *mpcE*, *mpcF*, *mpcD*, *mpcA* and *mpc* operon (collectively named *mpc* mutants) were generated in *B. abortus*. Each strain exhibited a significant growth defect on DOC at a concentration of only 0.005% (w/v) and the complemented mutant strains restored the wild-type phenotype (Fig. 1A). Furthermore, the MIC Test Strip stacking method on polymyxin B (Fig. 1B), a cationic peptide, indicated that the *mpc* mutants exhibited a slightly greater sensitivity (at 0.5 µg/mL) to polymyxin B compared to the WT strain (at 0.75 µg/mL). This finding suggests that the *mpc* mutants are unable to adapt to envelope stress.

**Figure 1.**
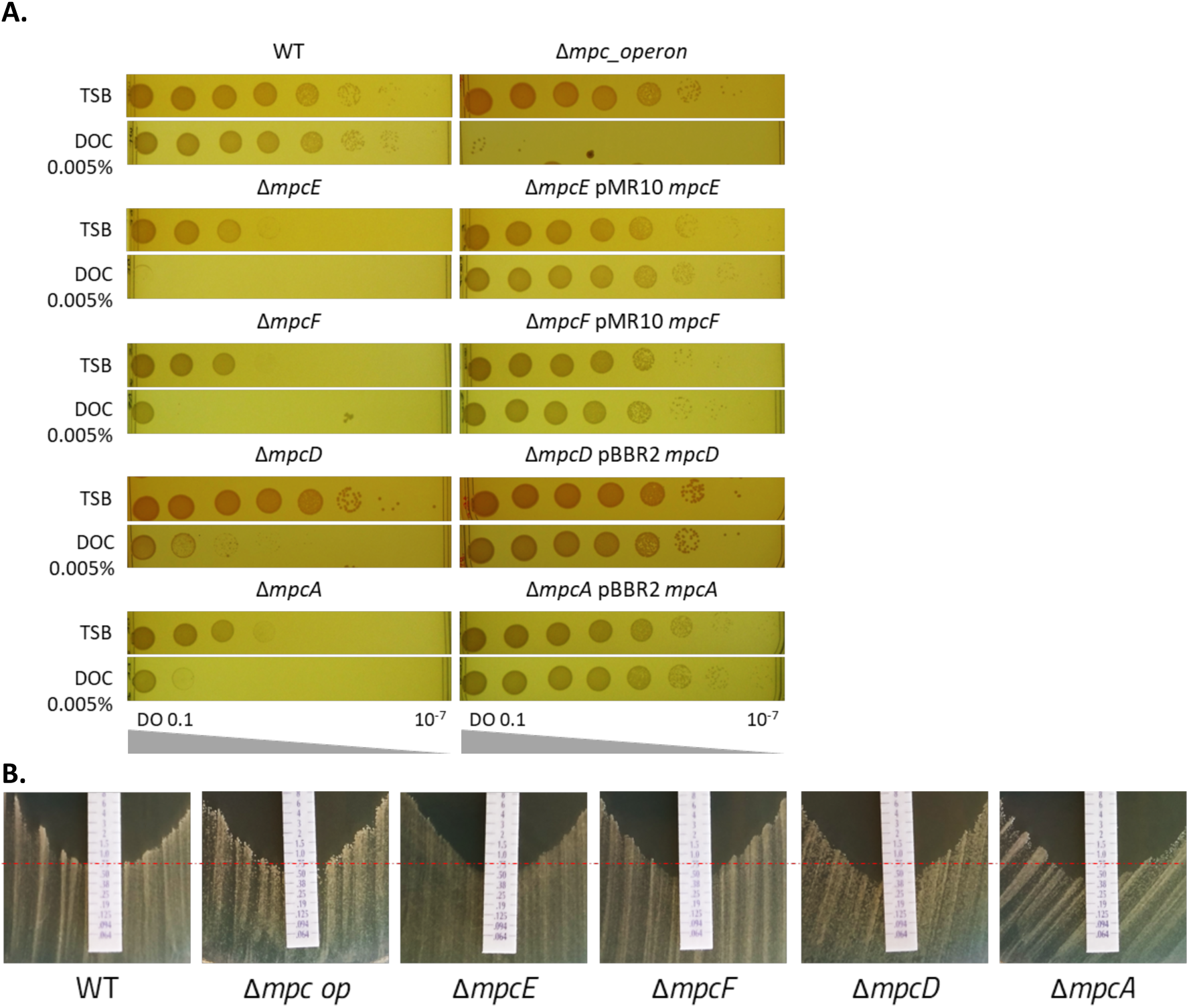
The *mpc* mutants exhibit a significant growth defect on DOC. **A.** A plating assay shows that *mpc* mutants are sensitive to DOC. The overnight cultures were normalized to 0.1 OD_600_ and serially diluted before being plated onto TSA with or without DOC. All *mpc* mutants exhibit a growth defect on DOC at 0.005%. The complemented strains exhibit a wild-type growth phenotype. All mutants except Δ*mpcD* also display a growth defect on TSA without DOC. **B.** MIC Test Strip stacking methods demonstrated the sensitivity to polymyxin B of *B. abortus* WT and the *mpc* mutants.

### The Mpc system is required for survival in macrophages

To test if envelope stress sensitivity of the *mpc* mutants correlates with a defect in virulence, J774.A1 macrophages were infected with these strains. The number of viable intracellular bacteria was evaluated with a colony-forming units (CFU) counting after gentamycin treatment, which kills extracellular bacteria. The CFU counts of the mutants were significantly lower at 2 and 5 hours post-infection (Fig. 2A). At 24 and 48 hours post-infection, the mutant strains remained at a lower level than the wild type (WT), but their intracellular growth was not affected (Fig. 2A and S2). To determine the ability of the bacteria to adhere and enter host cells, a double labelling was performed before and after cell permeabilization. The results showed that the ability of the *mpc* mutants to adhere and enter the macrophages was very similar to the WT strain (Fig. 2B). These results suggest that *mpc* mutants present survival defects in macrophages during the first hours of infection.

**Figure 2.**
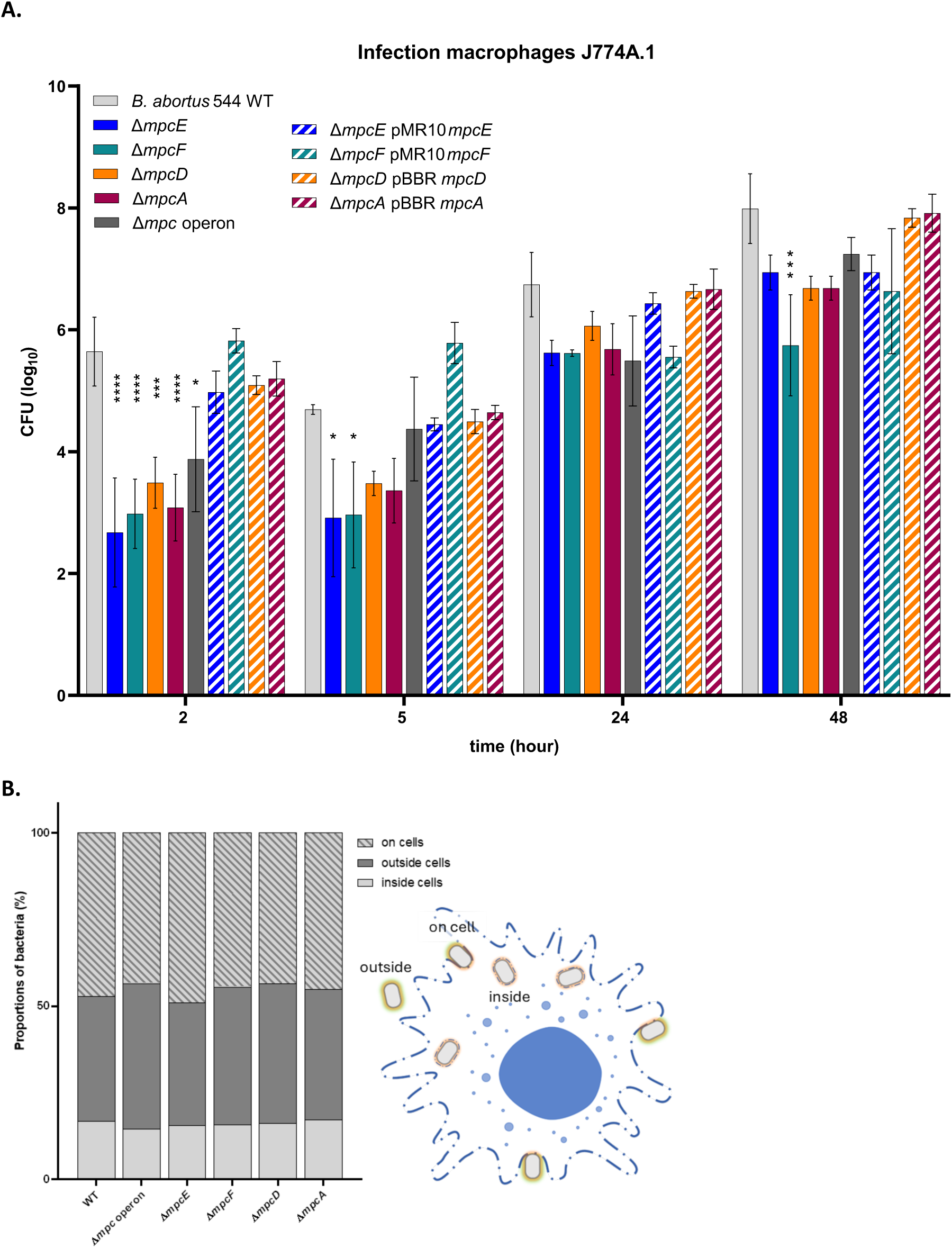
The *mpc* mutants are killed during early trafficking. **A.** Intracellular replication of WT, *mpc* mutants and the complemented strains were assessed by counting colony-forming units (CFU) at 2, 5, 24, and 48 hours post-infection of J774.A1 macrophages. The data represents the mean ± SD and were compiled from three independent replicates. Statistical significance between the results for a given strain and those for the WT was determined using a two-way ANOVA (not significant: p≥0.05 (not shown); *0.05<p<0.01, **0.01<p<0.001, ***0.001<p<0.0001, ****p< 0.0001). **B.** To identify the localization of bacteria inside, outside or on the macrophages, a double labelling technique was performed. After 2 hours post-infection, the cells were fixed with 4% PFA and the bacteria were labelled with a rabbit serum against *Brucella* (<u>Deghelt *et al*., 2014</u>). After the first set of labelling, the cell membrane was permeabilized with Triton X-100 0.1% and the bacteria were labelled a second time with a mouse monoclonal antibody against *Brucella abortus* lipopolysaccharide. Bacteria that were labelled only with the rabbit anti-*Brucella* serum are inside the cell, while those that were labelled with both antibodies are outside the cell. Bacteria adhering to the surface of the macrophages were found labelled with the mouse monoclonal antibodies on a part of the bacterial cell only. For each strain, the number of bacteria was counted by two individuals in 108 macrophages. The mean number of bacteria is presented as a percentage, depending on their location. The proportions of each localization type are unchanged for the mutant strains compared to the wild type, indicating that adherence and entry of the bacteria into the macrophages was not affected, while survival of the mutant was strongly impaired.

### The outer membrane lipid composition depends on Mpc

In *E. coli*, the Mla pathway is implicated in the retrograde transport of phospholipids between the outer and inner membranes (<u>Malinverni & Silhavy, 2009</u>; <u>Tang *et al*, 2021</u>; <u>Wotherspoon *et al*, 2024</u>). To investigate the possible function of Mpc pathway as a PLs transporter, outer membrane vesicles (OMVs) were isolated to characterize their PLs composition. Total lipid extraction was performed on OMVs and whole cells samples from *mpc* mutants and the WT. The thin layer chromatography (TLC) on lipid extractions showed that all *mpc* mutants have less amount of cardiolipin (CL) in OMVs compared to the WT (Fig. 3A and S3). Furthermore, there appeared to be no variations in PLs composition between the mutants and the WT in whole bacterial lysates. These findings suggested that the Mpc system may act as an PLs transport system.

**Figure 3.**
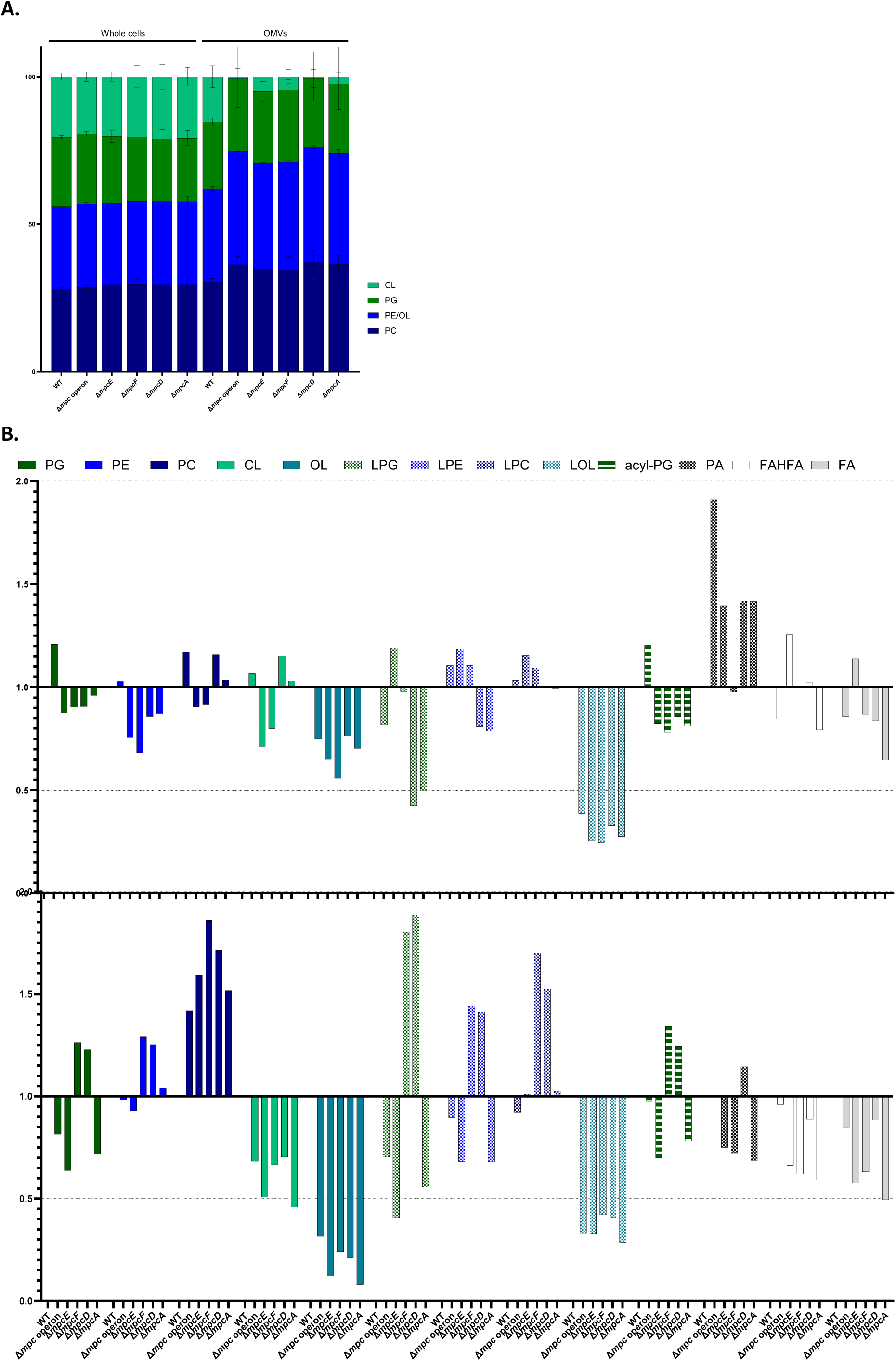
Lipid composition of whole cells compared to outer membrane vesicles. **A.** Quantification of bands of thin layer chromatography was performed using Quantity One software. The mean ± SD band intensity from three independent replicates was normalized to obtain the percentage composition of lipids. The stacked bars represent the percentages for each strain in both whole cells and outer membrane vesicles. PC: phosphatidylcholine, PE: phosphatidylethanolamine, OL: ornithine lipid, PG: phosphatidylglycerol, CL: cardiolipin. **B.** Bar plot of the ratio of the abundance of phospholipids identified by mass spectrometry-based lipidomic analysis was performed by comparison of the relative lipid concentration (µM) of *mpc* mutants to that of the WT strain. The whole cells samples are represented in top panel and the OMVs in bottom panel. PG: phosphatidylglycerol, PE: phosphatidylethanolamine, PC: phosphatidylcholine, OL: ornithine lipid, CL: cardiolipin, LPG: lyso-phosphatidylglycerol, LPE: lyso-phosphatidylethanolamine, LPC: lyso-phosphatidylcholine, LOL: lyso-ornithine lipid, acyl-PG: aminoacyl-phosphatidylglycerol, PA: Phosphatidate, FAHFA: fatty acid esters of hydroxy fatty acid, FA: fatty acid.

To obtain more information on the OMVs lipid composition, we analysed the total lipid composition using liquid-chromatography coupled to quadrupole-time-of-flight mass spectrometry, LC-MS (QTOF). The analysis revealed that the lipids composition of OMVs and whole cells are different but share commonalities in the changes associated with the *mpc* mutants. Significant differences were observed between the WT and *mpc* mutants in both whole cells and OMVs lipid extracts. These differences included a consistent decrease in the concentration and relative proportion of CL in OMVs lipid extracts and of ornithine lipids (OL) and lyso-ornithine lipids (LOL) in whole cells (Fig. 3B). The lipid extracts from all mutants OMVs have a profile that differ significantly from that of the WT, while the profiles of whole cells samples are more similar (Fig. S4). These data confirm that OMVs exhibit a significant reduction in CL in the *mpc* mutants and reveal that OL and LOL are also less abundant in whole cells of these mutants.

To ascertain whether the observed stress sensitivity phenotype is a consequence of the absence of CL, LOL or OL within the OM or their accumulation in the IM. The phenotype of a cardiolipin synthase (Cls) and ornithine lipid synthases (OlsA and OlsB) deletion mutants were subjected to analysis (Fig. 4A). The deletion mutants, which no longer possess CL, LOL or OL, did not exhibit any sensitivity in the presence of DOC (Fig. 4B) or in the infection model (Fig. 4C). This suggests that the absence of CL, LOL or OL is not sufficient to cause DOC sensitivity and attenuation in this cellular infection model. However, when combined with the *mpc* operon deletion, a significant increase in DOC sensitivity and attenuation in J774.A1 macrophages was observed in comparison to *mpc* mutants, only for the *cls* and *olsB* deletions. The deletion of *olsA* in the background of *mpc* operon deletion restores the WT phenotype in both conditions (Fig. 4B-C). This result suggests that the putative accumulation of LOL, induced by the absence of OlsA, could compensate the absence of Mpc system.

**Figure 4.**
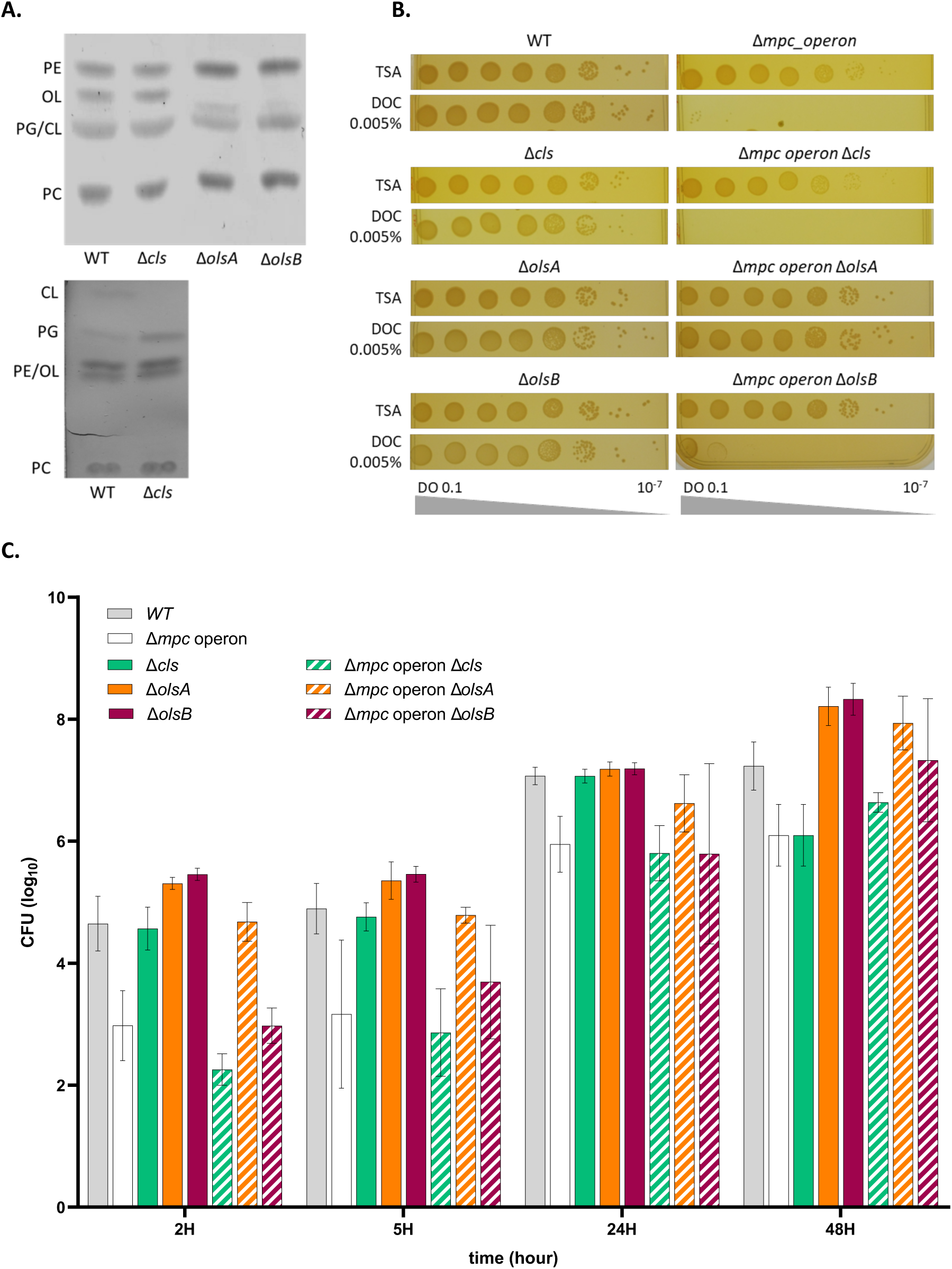
The sensitivity phenotype of *cls*, *olsA* and *olsB* deletion mutants. **A.** Thin layer chromatographiy (TLC) was performed on lipid extracts obtained from whole cells of lipid synthase mutants (1, WT; 2, Δ*cls; 3,* Δ*olsA; 4,* Δ*olsB*). On the top panel the lipids were separated by using a solution of TCM, MeOH and H_2_O_dd_ (14:6:1) and in the bottom panel of TCM, MeOH and AcOH (13:5:2). PC: phosphatidylcholine, PE: phosphatidylethanolamin, OL: ornithine lipids, PG: phosphatidylglycerol and CL: cardiolipin. **B.** The plating assay on *cls, olsA, olsB* and *asmA* mutants, *mpc* operon mutant and double mutants on DOC. All the mutants with deletion of *mpc* operon present the same sensitivity to DOC at 0.005%, exept for the double mutant with *olsA* which growth like the WT strain. The overnight cultures were normalized to 0.1 OD_600_ and serially diluted before being plated onto TSA with or without DOC. **C.** Intracellular replication of WT, *cls* mutant, *mpc* operon mutant and the double mutants were assessed by CFU at 2, 5, 24, and 48 hours post-infection of J774.A1 macrophages. The data represents the mean ± SD and were compiled from three independent replicates. Statistical significance between the results for a given strain and those for the WT was determined using a two-way ANOVA (not significant: p≥0.05 (not shown); *0.05<p<0.01, **0.01<p<0.001, ***0.001<p<0.0001, ****p< 0.0001).

### Mpc bridges inner to outer membrane

The *mpc* operon in *B. abortus* genome consists of only four genes, compared to six genes encoding for the Mla pathway in *E. coli*. Structural modelling was performed to determine how this simplified system could transport PLs between both membranes (Fig. 5A). The ABC transporter in IM is conserved, with the N-terminal domain of MpcE aligning with MlaB. The 100 first amino acids of MpcD aligns with MlaD. However, the MpcD (331 aa) is almost twice as long as MlaD (183 aa). The extended C-terminal domain of MpcD is a conserved feature among several Hyphomicrobiales, with the domain being even more extended in *Agrobacterium tumefaciens* and *Sinorhizobium meliloti*. The secondary structure prediction also exhibits conservation among Hyphomicrobiales, displaying a series of alpha-helices at in the C-terminal part of the protein (Fig. S6). Interestingly, the lipoprotein MpcA (encoded by the last gene of the *mpc* operon) shows homology with the PqiC protein. Additionally, there is no detectable homologue for MlaC in the *B. abortus* genome. The system was named the Mla-Pqi chimeric (Mpc) system as it encodes for a hybrid system between the Mla pathway and Pqi system (Fig. 6).

**Figure 5.**
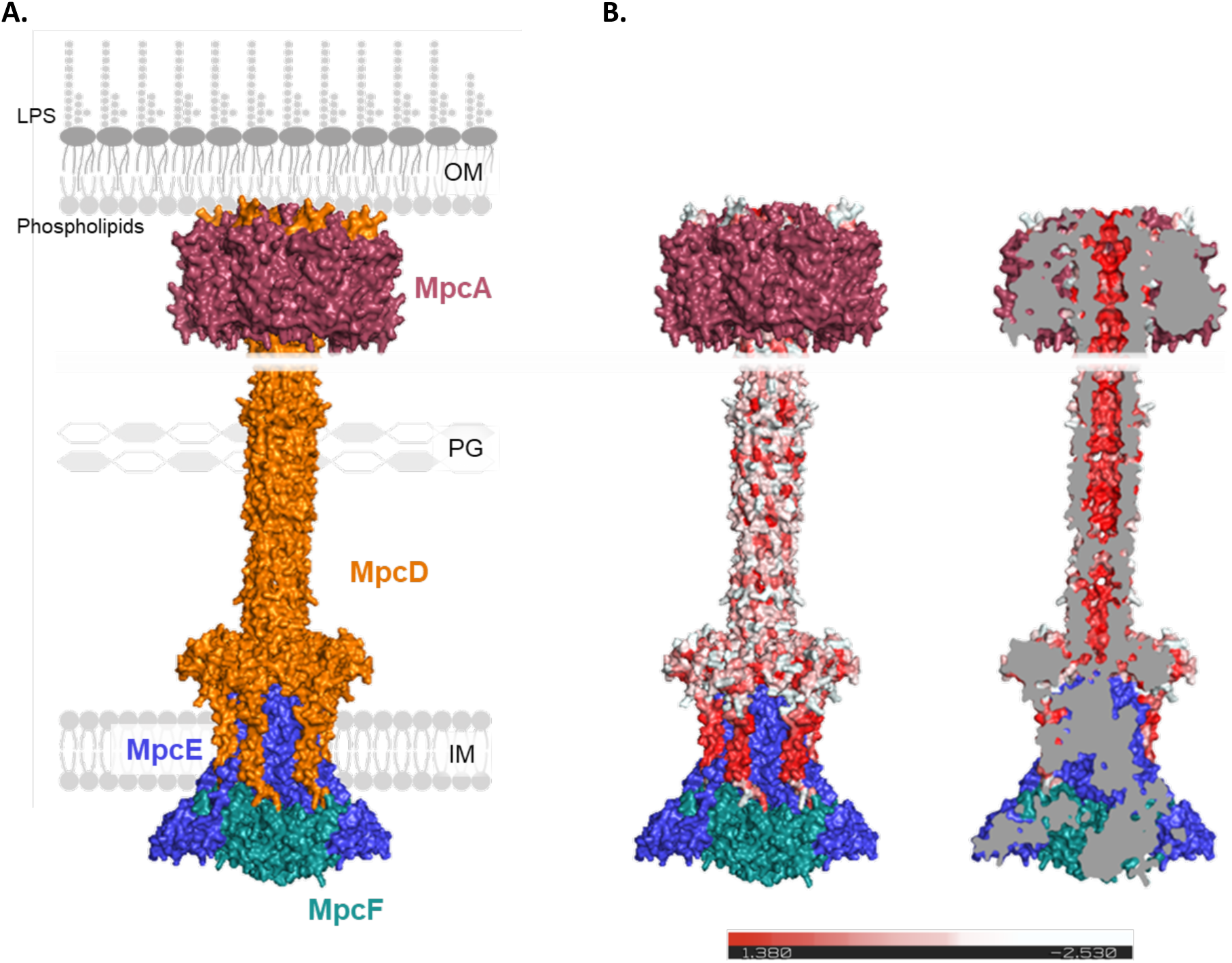
The multimer predictions of Mpc complex. **A.** Hydrophobicity (in red) of the full-length hexameric MpcD model was calculated using PyMol color_h command. The model has a length around ∼250 Å with transmembrane domain in N-terminal part. The tunnel of this model displays a hydrophobic interior, with a diameter of around ∼18 Å. In the cut-away view (right panel). **B.** Model of the interaction between Mpc proteins predicted with AlphaFold3 multimer predictions of biomolecular interactions (<u>Abramson *et al*., 2024</u>) of MpcE_2_F_2_D_6_ (bottom) and MpcD_6_A_8_ (top).

**Figure 6.**
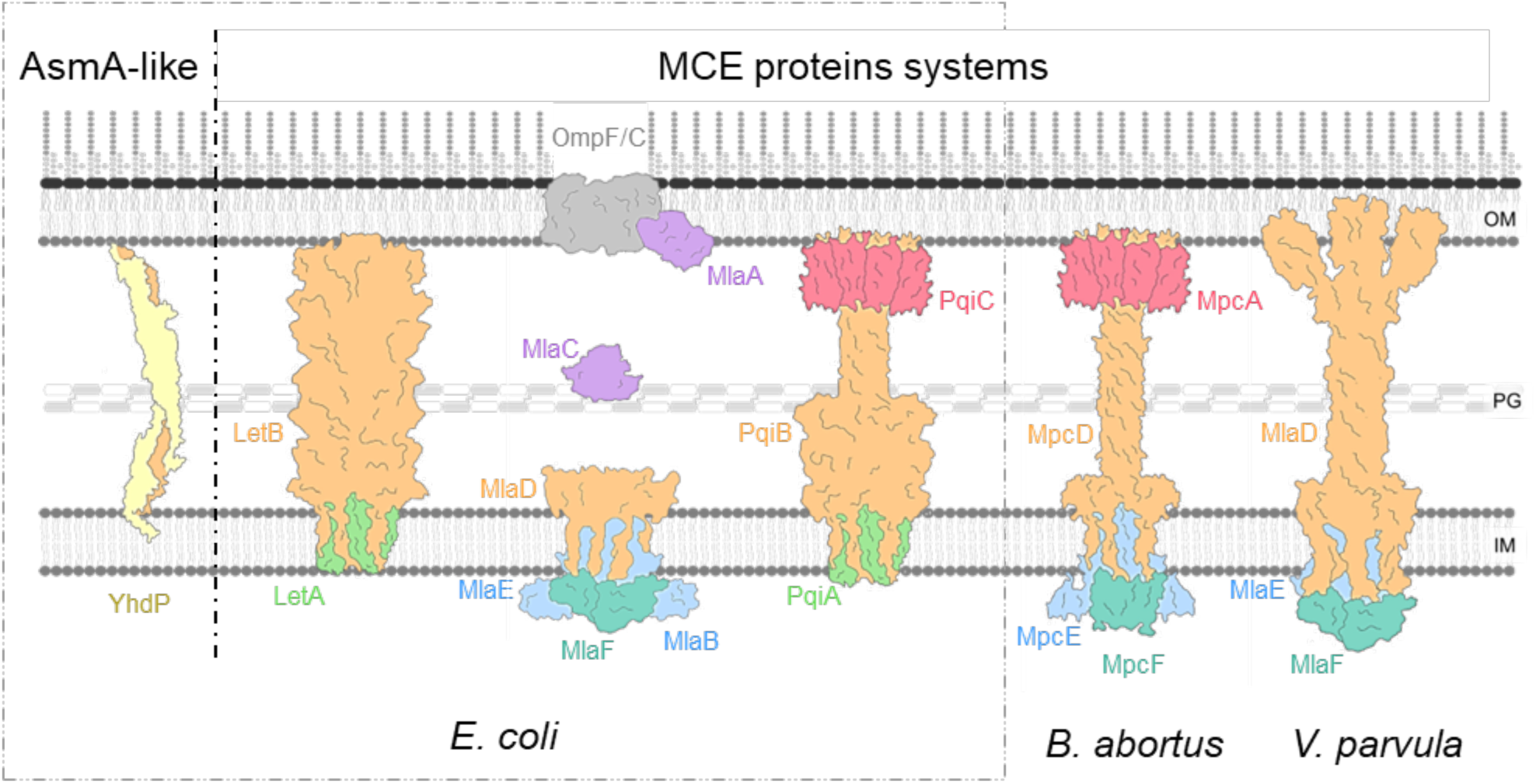
AsmA-like proteins and MCE proteins phospholipids transporters. Schematic representations of phospholipids transporters in *Escherichia coli*. AsmA-like proteins are represented by YhdP structure, from AlphaFold (<u>Cooper *et al*., 2023</u>). Let system, Mla pathway and Pqi system are comprising MCE proteins (orange). LetB (6V0C) structure was determined by Cryo-EM (<u>Isom *et al*.,</u> <u>2020</u>), LetA structure is not characterized. MlaEFDB (6XBD) structure was determined by Cryo-EM (<u>Coudray *et al*., 2020</u>), MlaC (5UWA) structure was determined by Cryo EM (<u>Ekiert *et al*., 2017</u>) and OmpF-MlaA (5NUO) structure was determined by X-rays diffraction (<u>Abellon-Ruiz *et al*, 2017</u>). Multi-MCE domain structure of PqiB (5UVN) was determined by Cryo-EM (<u>Ekiert *et al*., 2017</u>). PqiC (8Q2C) X-rays diffraction structure was obtained and the structure of PqiABC was predicted by AlphaFold-Multimer (<u>Cooper *et al*., 2024</u>). The structure of MpcEFDA system in *Brucella abortus*, was predicted by AlphaFold-Multimer in this study. For the Mla system homologs in *Veillonella parvula*, the structure of MlaD was predicted by AlphaFold2 and the homologs of MlaE and MlaF were identified as conserved (<u>Grasekamp *et al*., 2023</u>).

The AlphaFold 3 multimer predictions of biomolecular interactions (<u>Abramson *et al*, 2024</u>) of Mpc proteins and some oleic acids as model lipids were conducted. As the number of monomers in the MlaE_2_F_2_D_6_B_2_ and PqiA_2_B_6_C_8_ complexes is known (<u>Cooper *et al*., 2024</u>; <u>Ekiert *et al*., 2017</u>; <u>Malinverni &</u> <u>Silhavy, 2009</u>; <u>Nakayama & Zhang-Akiyama, 2017</u>), we used the same monomer number for the Mpc complex prediction, MpcE_2_F_2_D_6_A_8_. The prediction for the entire complex was unsuccessful due to the coiled-coil alpha-helices of the MpcD_6_ channel being folded over themselves. Subsequently, we proceeded to predict the interactions of MpcD_6_ with MpcA_8_ and, in another instance, with MpcE_2_F_2_ in the presence of oleic acids that could potentially mimic interactions with lipids (Fig. S7). The MpcE_2_F_2_D_6_ AlphaFold3-predicted structure exhibited high confidence for the MCE, transmembrane and cytoplasmic domains, whereas the alpha-helices series displayed low confidence. However, the prediction confidence of the coiled-coil alpha-helices prediction is enhanced in the presence of oleic acids (Fig. S7B). This suggests that the hydrophobic side chain inside the tunnel formed by the alpha-helices decreased confidence in the model. The MpcD_6_A_8_ AlphaFold3-predicted structure exhibits high confidence in the octameric form of MpcA and lower confidence in the hexameric form of MpcD than in the MpcE_2_F_2_D_6_ structure prediction (Fig. S7A). A fusion of the two AlphaFold3-predicted structures is illustrated in Figure 5A. The complex size is sufficient to span the entire periplasmic space. Additionally, the predicted hexameric structure of MpcD reveals an internal hydrophobic channel (Fig. 5B) that may facilitate the transport of hydrophobic compounds, such as lipids. To confirm the interaction of Mpc proteins, a pull-down assay was conducted on MpcA fused with a Strep-tag, followed by analysis through liquid chromatography coupled with tandem mass spectrometry (LC-MS/MS). Among all the proteins in the proteome, we specifically identified three proteins, MpcE, MpcF and MpcD, as co-precipitating with MpcA. (Fig. 7). These data indicate that MpcEFDA form a stable complex, from the cytoplasm to the OM.

**Figure 7.**
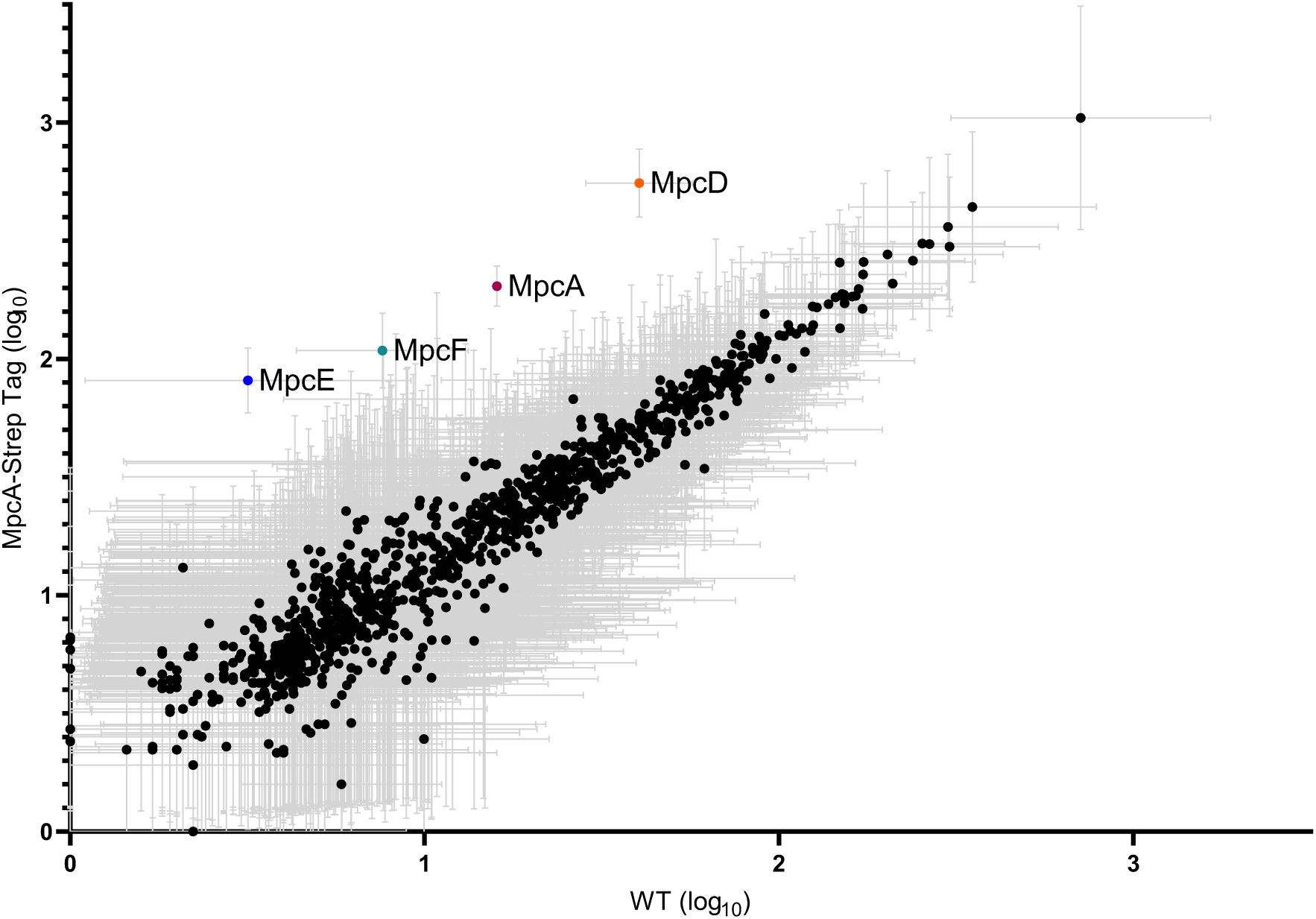
MpcEFDA is a stable complex. Proteins co-purified with MpcA-Strep Tag tagged were identified by MS. The data represents the mean ± SD of Total Spectrum Count in log_10_ for each identified protein (1508) from three independent replicates. The Total Spectrum Count obtained from the pull-down assay on MpcA-Strep Tag is related to the pull-down assay on WT (negative control).

## Discussion

The present study characterises a newly identified complex spanning the periplasmic space, namely the Mpc (Mla-Pqi chimeric) system, connecting IM to OM in *B. abortus*. All mutants generated in this system were unable to grow in the presence of DOC (Fig. 1A), a compound known to affect bacterial membrane integrity (<u>Urdaneta & Casadesus, 2017</u>). It is noteworthy that these mutants also exhibit sensitivity to polymyxin B (Fig. 1B), indicating that the Mpc system may be crucial for survival in the presence of cationic peptides generated by immune cells (<u>Sawyer *et al*, 1988</u>). Conversely, it is well-established that wild-type *Brucella* strains exhibit high resistance to cationic peptides (<u>Martínez de Tejada *et al*, 1995</u>). This suggests that these bacteria encounter cationic peptides during the natural infection, or other envelope stress conditions unidentified so far. Given the inability of the *mpc* mutants to survive to the initial stages of infection in macrophages (Fig. 2), we propose that *B. abortus* requires this system to survive to the aggression of its envelope at the beginning of its intracellular trafficking. At this stage of the trafficking process, *B. abortus* resides within endosomal *Brucella*- containing vacuoles (eBCVs), where bactericidal conditions are particularly important (<u>Celli *et al*, 2003</u>; <u>Porte *et al*, 1999</u>). These eBCVs are characterised by an acidic pH (<u>Porte *et al*., 1999</u>) and the presence of markers of late endosomes, including LAMP1 and Rab7 (<u>Celli *et al*., 2003</u>; <u>Starr *et al*, 2008</u>). Furthermore, it has been demonstrated that *B. abortus* growth is arrested in these compartments (<u>Deghelt *et al*, 2014</u>), which may be necessary to enable survival in the presence of adverse conditions such as starvation, acidic pH and envelope stress encountered in eBCVs. The necessity of the Mpc system in other infection models is corroborated by the essentiality of the *mpc* genes in Tn-seq analyses conducted on macrophages and mice infection models (<u>Potemberg *et al*, 2022</u>; <u>Sternon *et al*.,</u> <u>2018</u>). The sensitivity of the *mpc* mutants to the infection conditions correlates with an alteration in the composition of the lipids present in the OMVs. Given the homology of the Mpc system to the Mla and Pqi systems, which are involved in the transport of lipids between the IM and OM (<u>Coudray *et al*.,</u> <u>2020</u>; <u>Malinverni & Silhavy, 2009</u>; <u>Nakayama & Zhang-Akiyama, 2017</u>; <u>Tang *et al*., 2021</u>), it is very likely that the Mpc system mediates the exchange of lipids between IM and OM in *B. abortus*. This hypothesis suggests that the lipid composition of the OM may be a crucial factor in enabling *B. abortus* to respond effectively to adverse conditions in eBCVs. Furthermore, alterations of the OM composition may also provide an explanation for the recurrent isolation of *asmA*-like mutants in Tn-seq screens with *B. abortus* or *Brucella melitensis* in macrophages and mouse models of infection (<u>Potemberg *et al*.,</u> <u>2022</u>; <u>Sternon *et al*., 2018</u>). It is noteworthy that the *asmA* mutant has also been isolated in the DOC sensitivity Tn-seq (Table 1), which indicates that the AsmA-like protein could have a role in maintaining the integrity of the OM. This is consistent with the proposed AsmA-like function of bidirectional diffusive flow of phospholipids from the IM to the OM (<u>Kumar & Ruiz, 2023</u>) being conserved in *B. abortus*. The double deletion of *asmA* in the *mpc-operon* mutant did not result in a synthetic phenotype in response to DOC (Fig. S5A-B), suggesting that AsmA and Mpc may be involved in similar but non-redundant physiological functions related to lipid transport. Collectively, these results indicate that the composition of the OM is a critical determinant of host-pathogen interactions. Further investigations are necessary to gain a deeper understanding of OM properties, including its composition, structure, biogenesis and its role in responding to various stresses encountered during the infectious cycle.

Our results, as illustrated in Figure 3, indicate a decrease in the proportion of CL in OMVs and a general decrease in OL and LOL in *mpc* mutants. With this result, it is not possible to determine the direction of lipid transport. Indeed, the absence of CL in OMVs in the *mpc* mutants could be attributed to either the lack of anterograde (IM to OM) CL transport or to the accumulation of other PLs in OM in the absence of retrograde (OM to IM) transport of PLs, which would dilute CL in the OM. In light of the observed defect in OM integrity in the *mpc* mutants, it can be postulated that the observed decrease in OL and LOL may result from a rescue phenomenon. Indeed, if the PL transport is imbalanced in the absence of the Mpc system, it is plausible that the bacteria adjust the pools of PLs to compensate for the absence of one of the PLs transporters, resulting in a decrease in the relative concentration of OL and LOL in the envelope. Moreover, our results demonstrate that the observed sensitivity phenotype is effectively induced by the absence of the Mpc system, rather than by the lower abundance of CL, OL or LOL in the OM (Fig. 3). The phenotype of the *cls* mutant, which encodes a non-redundant CL synthase in the tested condition (Fig. 4A), is similar to the wild-type strain in terms of DOC sensitivity and survival at short times post-infection (Fig. 4B-C)(<u>Palacios-Chaves *et al*, 2011</u>). The *olsA* and *olsB* deletion mutants, which encode ornithine lipid biosynthesis, exhibited identical results to the WT in terms of DOC sensitivity and survival at short times post-infection, in accordance with previous observations (<u>Palacios-Chaves *et al*., 2011</u>). The conversion of ornithine to OL occurs in two stages. The initial step is catalysed by OlsB, which adds a fatty acyl group to ornithine to form LOL. The second step is the addition of a fatty acyl group to the first fatty acid, transforming LOL to OL through the action of OlsA (<u>Gao *et al*, 2004</u>). However, the deletion of *olsA* in the *mpc* operon background allows a restoration of the wild-type phenotype (Fig. 4B-C). This suggests that the accumulation of LOL (<u>Gao *et*</u> *<u>al.</u>*<u>, 2004</u>) may have an impact on the DOC sensitivity and survival at short times post-infection only when the Mpc system is absent. It has been demonstrated that the accumulation of lyso-PLs is involved in the stress resistance and virulence of bacteria (<u>Zheng *et al*, 2017</u>). We cannot exclude that other modifications to the lipid composition of the membranes may also affect the survival rate at short times post-infection. Collectively, these findings indicate that the absence of the Mpc system may have a significant impact on the lipid composition of membranes, due to a defect in lipid transport.

The proposed structure of Mpc forms a channel between IM and OM (Fig. 6A). The size of the complex, from the N-terminus of MpcA to MpcE is consistent with the measure of the periplasm width evaluated from CryoEM pictures (<u>Godessart *et al*., 2021</u>). The most notable aspect of the structure is the interaction between the MpcA lipoprotein and the hexameric coiled coil of MpcD, the latter forming an entirely hydrophobic channel. We propose that the octameric form of MpcA engages in a cap-like interaction with MpcD, similarly to the proposed role of PqiC in anchoring the PqiB periplasmic needle and limiting its flexibility (<u>Cooper *et al*., 2024</u>). This hypothesis is corroborated by the results of the pull-down assay (Fig. 5), which demonstrated that Mpc interactions are strong and form a stable complex. At the evolutionary level, it is not excluded that Mpc systems may have been ancestral, leading to diversification in different systems (Mla and Pqi). It is indeed likely that intermembrane complexes with various architecture based on one or several MCE domains and a hexameric alpha-helical coiled coil are relatively ancient since they are also found in *Veillonella parvula* but with a beta-barrel inserted in the OM (Fig. 6) (<u>Grasekamp *et al*., 2023</u>). The complete hydrophobic nature of the channel formed by hexameric MpcD is consistent with the channelling of acyl chains of the PLs. However, the amphipathic nature of the PLs must be accommodated in such a hydrophobic tunnel, a process that will be elucidated by future investigation. Homologs of the Mpc system, with a predicted structure of elongated alpha-helical C-terminal domain for MpcD, are also found in many Hyphomicrobiales, but with a large variation in the size of the channel (Fig. S6). This large variation may reflect different periplasmic width or different angles of the MpcD relative to the IM and OM. It can therefore be surmised that the Mpc system described here is a conserved system, at least in the Hyphomicrobiales, whose function would require further investigation in several other bacteria of interest for basic knowledge, health and biotechnology.

## Materials

### Bacterial strains and media

*Brucella abortus* 544 Nal^R^ strain (referred to the WT in this study) and its derivates were cultivated in TSB-rich medium (Difco^TM^ Tryptic Soy Broth) at 37°C. All *Escherichia coli* strains were grown in LB Broth Base (Lennox formulation) at 37°C. All the strains used in this study are listed in Table S1. Antibiotics were added, when it is necessary, at the following concentrations: kanamycin (10 µg/mL or 50 µg/mL for genomic or plasmidic resistance), nalidixic acid (25 µg/mL) and gentamicin (20 µg/mL) ampicillin (100 µg/mL).

### Strains construction

All bacterial strains, plasmids, open reading frames and primers used in this study are listed in the supplementary tables 1–4, respectively. For deletion strains, approximately 500 bp upstream and downstream of the coding sequence of interest were amplified from the purified *B. abortus* 544 gDNA by PCR using Q5® High-Fidelity DNA Polymerase (New England Biolabs). The fragments were assembled by overlapping PCR. For the MpcA-StrepTag strain, 500 bp upstream and downstream of the C-terminal region of *mpcA* gene and the StrepTag sequence (AWSHPQFEK) with a linker (GGGSGGS) were amplified by PCR. The fragments were assembled by overlapping PCR. For complemented strains the genes of interest were amplified. The resulting amplicons were purified from agarose gels and digested as the destination vector with the corresponding restriction enzymes (New England Biolabs) and ligated (T4 DNA ligase Promega) overnight at 20°C. The ligation products were transformed in *E. coli* DH10B, and clones were screened by PCR using GoTaq® DNA polymerase (Promega). Selected plasmids were checked by sequencing and transformed in *E. coli* S17-1 or MFDpir strain to allow conjugation to *B. abortus* 544 Nal^R^ strain. Fusion and deletion strains were made by allelic exchange on the chromosome using a non-replicative plasmid, pNPTS138 (<u>Deghelt *et al*., 2014</u>). Complemented strains were made using a replicative plasmid (pBBR2MCS or pMR10) and the genes of interest were under the control of the *E. coli lacZ* promoter or their endogenous promoter, respectively.

### Transposon sequencing assay

One milliliter of an overnight culture of *B. abortus* 544 Nal^R^ strain was mixed with 50 µL of an overnight culture of the conjugative *E. coli* MFDpir strain carrying the pXMCS-2 mini-Tn*5* Kan^R^ plasmid (<u>Sternon</u> *<u>et al.</u>*<u>, 2018</u>). The plasmid facilitated the straightforward generation of a high number of transposon mutants due to the hyperactive Tn*5* transposase encoded by the plasmid. The mating mixture was incubated overnight at room temperature (RT) on TSB agar plates supplemented with *meso-*2,6-Diaminopimelic acid (300 µM). The resulting *B. abortus* mini-Tn*5* libraries were selected on TSB agar plates supplemented with kanamycin. The mutant library with envelope stress condition, the medium was supplemented with sterile 0.015% of sodium deoxycholate (w/v). Mini-Tn*5* mutagenesis generates insertion of the transposon at only one locus per genome, as demonstrated previously for Brucella (<u>Lestrate *et al*, 2000</u>; <u>Sternon *et al*., 2018</u>).

### Analysis of essential genes for growth

Genomic DNA was extracted from each transposon library using standard techniques and prepared for sequencing of the mini-transposon library. *B. abortus* Tn mutants from each plate were collected, mixed in 2% SDS, and killed by heating (1 h, 80°C). The lysate was incubated for 3 days at 37°C under constant agitation in TENa buffer (pH = 8.0, Tris 50 mM, EDTA 50 mM, 100 mM NaCl) which was complemented with 0.1 mg/mL Proteinase K (Merck) and 0.5% SDS. Subsequently, an equal volume of isopropanol was added to precipitate the gDNA, which was then washed with 70% ethanol. The gDNA was resuspended in deionised water and the regions flanking the mini-Tn*5* were sequenced (Fasteris company, Geneva, Switzerland). In order to map the insertion sites of mini-Tn*5*, the libraries were sequenced on an Illumina HiSeq with a primer hybridised at the border of the transposon, with its 3’ end pointing towards the flanking gDNA. Raw reads were filtering using cutadapt algorithm and then mapped on *B. abortus* 544 genome using Burrows-Wheeler Alignment (BWA) algorithm (114,039 mapped reads in Control condition and 134,058 mapped reds in DOC condition). The read counts were determined using samtools (<u>Coppine *et al*, 2020</u>).

In the control condition, there was an average of one unique insertion site every 2.62 bp, and in the envelope stress condition, every 2.48 bp, saturating the *B. abortus* genome in both conditions. The mapping analysis (<u>Coppine *et al*., 2020</u>) yielded an insertion number per bp for 80% of each open reading frame (ORF - excluding the first and last 10% of the ORF). Essential genes were identified based on the insertion number per bp. An open reading frame (ORF) is considered essential if the insertion number per bp is below 0.1, which corresponds to one insertion every 10 bp (Fig. S1). Out of the predicted 3,390 ORF in the B. abortus genome, 551 have been identified as essential for growth in rich medium, which correspond to 16.2% of the predicted genes. To enable a quantitative analysis of each ORF in different conditions, we calculated a transposon insertion frequency (TnIF) parameter. This frequency corresponds to the logarithm in base 10 of the mapped read numbers (*ES.Reads*) for the central 80% of the ORF [log_10_(*ES.Reads*+1)]. To identify genes required for growth in the presence of DOC, when the envelope is destabilised, we compared the TnIF of the stress condition to the control condition to obtain a ΔTnIF for each ORF (ΔTnIF = TnIF_DOC_ - TnIF_ctrl_). Thus, ORF with a negative ΔTnIF value correspond to mutants with a DOC-sensitivity phenotype (Table 1).

### Envelope stress growth tests

Overnight culture of *B. abortus* 544 and derivate strains are normalized at OD_600_ 0.1 and dilute in tenfold increments. Fifteen µL of each dilution are spotted on TSA plate containing or not sodium deoxycholate at 0.005% (w/v).

The MIC Test Strip (Liofilchem) is a quantitative assay used to determine the Minimum Inhibitory Concentration (MIC) of a microorganism. This is achieved by placing a strip comprising a gradient of Polymyxin B concentrations on an agar plate inoculated with *B. abortus* 544 and derivate strains.

### J774A.1 macrophage culture and infection

J774A.1 (ATCC) macrophages were cultured in Dulbecco’s Modified Eagle Medium (DMEM), GlutaMAX™ supplement medium (Gibco) supplemented with 10% heat inactivated foetal bovine serum (FBS, Gibco) at 37°C in a 5% CO_2_ atmosphere. The day before infection, cells were inoculated at 10^5^ cells/mL in 24 wells plate (0.5 mL per well). Overnight culture of *Brucella* was prepared in DMEM at a multiplicity of infection (MOI) of 50 bacteria per cell. The suspension of bacteria was place on the cell and centrifuged at 169 rcf for 10 min at 4°C. Infected cells were incubated for 1 hour at 37°C in a 5% CO_2_ atmosphere. Subsequently, the culture medium was removed and replaced with fresh medium supplemented with 50 µg/mL gentamicin to kill extracellular bacteria. After 1 hour, the medium was changed to DMEM with 10% FBS with 10 µg/mL gentamicin.

### CFU counting

At 2 h, 5 h, 24 h and 48 h post-infection, cells were washed twice with phosphate-buffered saline (PBS), and macrophages were lysed using PBS–0.1% Triton X-100 for 10 min at 37°C and flushed to recover bacteria. Twenty µL of dilutions bacteria were plated onto TSB agar plate and the colony forming unit (CFU) were counted after incubation at 37°C.

### Immunolabeling of infected hosts cells

Labelling of intra-extra-bacteria in host cell, was carried out like previously reported (<u>Deghelt *et al*.,</u> <u>2014</u>) with minor modifications. Briefly, the cells were inoculated at 3*10^5^ cells/mL in a 24-well plate with coverslips. At 2h post-infection at MOI 1000, cells on the coverslips, were washed three times with PBS and fixed with 4% paraformaldehyde (PFA, ThermoFisher) at RT during 30 min. Coverslips were wash twice with PBS, and first label during 45 min at RT with a homemade rabbit serum against heat kill Brucella, diluted 2,000 times in PBS with 3% BSA. Subsequently the permeabilization with 0.1% Triton-X-100 in 3% BSA for 10 min, the coverslips were incubated for 45 min at RT with anti-lipopolysaccharide A76-12G12 monoclonal antibody (<u>Cloeckaert *et al*, 1993</u>) undiluted in 3% BSA, washed three times with PBS and incubated with secondary antibodies solution (each diluted 500 times) containing a goat anti-IgG mouse Alexa 514 (Sigma Aldrich) and an anti-IgG rabbit Pacific blue (Sigma Aldrich), in PBS with 3% BSA and 0.1% Triton-X-100, for 30 min at RT. The coverslips were washed three times with PBS, then once with demineralised water and mounted with Fluoromount-G™ Mounting Medium (Invitrogen).

### Microscopy and analysis

*Brucella* strains and infected J774A.1 cells were observed with a Nikon Eclipse Ti2 equipped with a phase-contrast objective Plan Apo λ DM100XK 1.45/ 0.13 PH3 and a Hamamatsu C13440-20CU ORCA-FLASH 4.0. Fluorescence images were analysed using FIJI v.2.1.0, a distribution of ImageJ (<u>Schindelin *et al*, 2012</u>). Look-up tables (LUT) were adjusted to the best signal–background ratio.

### Isolation of *Brucella* OMVs

Six 48-hour cultures of 800 mL were inactivated with 0.5% phenol and centrifuged at 8200 rcf at RT for 20 min at 4°C. The pellet was conserved, and the supernatant was concentrated using the Pellicon® tangential flow filtration system, with a membrane of 100.000 Dalton exclusion limit. The concentrated supernatant was centrifuged at 8200 rcf, 4°C for 20 min to discard the sediment. Then, to aggregate the OMVs, the supernatant was frozen at −20°C. The defrosted supernatant was ultra-centrifuged at 47.000 rcf, 4°C for 3 hours. The pellet, the isolated OMVs, was resuspended in deionized water and dialyzed at 4°C with deionized water for 3 days and finally frozen at −80°C. Both the culture pellet and isolated OMVs suspension were lyophilised by freeze dryer (TELSTAR CRYODOS 50).

### Lipidome analysis

Total lipids were extracted as described by Bligh and Dyer (<u>Bligh & Dyer, 1959</u>; <u>Daniels *et al*, 1993</u>), briefly, 500 µL of resuspended bacteria or OMVs (20 mg of lyophilised samples) in deionised water were vortexed for 3 hours with 650 µL of chloroform and 1300 µL of methanol. 650 µL each of chloroform and deionised water were add and mixed for 30 minutes. After short centrifugation to separate aqueous and organic phases, the organic phase was kept and evaporated with a flow of nitrogen.

Thin layer chromatography (TLC) was then performed on the lipid extractions to separate the phospholipids into different bands (<u>Ando & Saito, 1987</u>). The lipids extracts were analysed on silica gel 60 high-performance thin layer chromatography plates (Merck Chemicals) and chromatography was performed in the solution composed of chloroform (TCM), methanol (MeOH) and demineralised water (H_2_O_dd_)(14:6:1 respectively v:v). or TCM, MeOH and acid acetic glacial (AcOH) (13:5:2 respectively v:v). Plates were developed with a solution of CuSO_4_ (10%, w/v) in acid phosphoric (8%, v/v) at 180°C. The lipids were separated in 4 bands (1) phosphatidylcholine (PC), (2) phosphatidylethanolamine and ornithine lipid (PE/OL), (3) phosphatidylglycerol (PG), and (4) cardiolipin (CL). The intensity of each band was measured with GS-800™ densitometer (BioRad) and Quantity One – 4.6.6 software to enable comparison between samples.

Complementary to TLC analysis, a mass spectrometry-based lipidomic analysis was performed using ultra high-performance liquid-chromatography coupled to quadrupole-time-of-flight mass spectrometry, (LC-MS (QTOF)) as previously described (<u>Fernández-García *et al*, 2023</u>), with minor modifications. Briefly, lipid residues obtained by Bligh and Dyer for whole bacteria or OMVs were resuspended in 300 µL of methanol:chloroform containing 25 mg/L of d17:0 sphinganine (IS1). Reextraction of lipids was performed by vortexing at room temperature RT for 20 min. Lipid extracts were transferred to individual LC-MS vials, where 20 µL of SPLASH Lipidomix® lipid standard mixture (IS2, Avanti Polar Lipids, CA, USA) were added to each vial. Then, extracts were evaporated to dryness using a vacuum concentrator at 37°C. Residues were reextracted in 100 µL of methanol:chloroform (2:1, v/v) by thorough vortexing for 30 min at RT. Sample analysis was performed using an Agilent 1290 HPLC system coupled to an Agilent 6545 QTOF mass analyser (both from Agilent Technologies, Santa Clara, CA, USA), equipped with an ESI ionization source. Two run sequences of sample extracts corresponding to positive (ESI+) and negative (ESI-) ionization modes were performed. 2 (ESI+) or 5 μL (ESI-) were injected into an Agilent InfinityLab Poroshell 120 EC-C18 column (3.0 mm × 100 mm, 2.7 μm; Agilent Technologies, Santa Clara, CA, USA) equipped with an Agilent InfinityLab Poroshell 120 EC- C18 guard column (3.0 mm × 5 mm, 2.7 μm; Agilent Technologies, Santa Clara, CA, USA), operating at 50°C. A mobile phase flow rate of 0.6 mL/min was maintained throughout the chromatographic gradient. Firstly, 70% B was held until 1 min. Secondly, 86% B was achieved at 3.5 min and held until 10 min. Next, 100% B was achieved at 11 min and held until 17 min, followed by 2 min of re-equilibration time at 70% B. The total method runtime was 19 min. Metabolites were ionized using an ESI source with a nebulizer at 50 psi, a drying gas temperature of 200°C a drying gas flow rate of 10 L/min, a sheath gas temperature of 300°C, and a sheath gas flow rate of 12 L/min. In both polarity modes (ESI+ and ESI−), the capillary, fragmentor, skimmer, and octupole radiofrequency voltages were set to 3500, 150, 65, and 750 V, respectively. Firstly, full MS was selected as the data acquisition mode at an acquisition rate of 3.5 spectra/s over a mass range of m/z 50 to 3000 for ESI-. Next, two iterative auto MS/MS analyses (under both polarity modes) were conducted under identical chromatographic conditions, to provide with MS/MS data allowing high-confidence lipid annotation. MS/MS spectra were systematically acquired via iterative MS/MS mode over subsequent injections from a sample pool. Collision energy was set to 20 eV for ESI+, and 40 eV for ESI-. Lipids were annotated by using Agilent MassHunter Lipid Annotator (v. 1.0) followed by manual curation of annotations using Agilent MassHunter Qualitative Analysis (v. 10.0) considering lipid subclass diagnostic ions, adduct pattern, and elution order (<u>Köfeler *et al*, 2021</u>). OL and LOL were manually annotated using Agilent MassHunter Qualitative Analysis in accordance with reported fragmentation spectra (<u>Zhang *et al*, 2011</u>). The curated annotation data matrix was used as input for targeted compound integration using Agilent MassHunter Profinder (v. 10.0).

### Homology and structures predictions

Mpc proteins of *B. abortus* and *E. coli* Mla proteins were searched using standard protein BLAST with the DELTA-BLAST (Domain Enhanced Lookup Time Accelerated BLAST)(<u>Boratyn *et al*, 2012</u>). Secondary structures of homologues of MpcD in Hyphomicrobiales were predicted with NetSurfP-2.0 (<u>Klausen *et*</u> *<u>al</u>*<u>, 2019</u>). The interaction between Mpc proteins, MpcE_2_F_2_D_6_ and MpcD_6_A_8_, were predicted using AlphaFold3 multimer predictions of biomolecular interactions (<u>Abramson *et al*., 2024</u>). Resulting predictions were visualised and analysed in The PyMOL Molecular Graphics System, Version 2.4.1 Schrödinger, LLC.

### Pull-down assay

*B. abortus* cultures of 200 mL (0.8 OD_600_) were harvested by centrifugation, at 4500 rcf at 4°C for 20 min, then washed three times with PBS. The pellet was resuspended in 2 mL of DDM lysis buffer (CellLytic™ B cell lysis reagent (Sigma), 10 mM MgCl_2_, 0.05% Ready-Lyse lysozyme (v/v), 0.25 mg/mL DNAse I, 0.5% DDM (w/v), cOmplete™ EDTA free protease inhibitor cocktail) incubate 30 min at RT. The lysate was sonicated on ice by applying 6 bursts of 10 sec. at 60% amplitude and centrifuged at 12.500 rcf at 4°C for 15 min. Collect the supernatant and add 1 mg/mL avidin and incubate 30 min at 4°C a-on wheel then centrifuges for 5 min at 6000 rcf. The fusion protein MpcA-StrepTag was purified using the MagStrep “type 3” XT bead (Iba) in accordance with the manufacturer’s instructions.

The proteins recover in the elution are treated using Filter-aided sample preparation (FASP) to generate tryptic peptides. 40 µg of protein adjusted in 150 µl of buffer UA (8 M urea in 0.1 M Tris-HCl pH = 8,5) are filtrated on Microcon-30kDa (Millipore, MRCFOR030), previously rinsed with formic acid 1 %(v/v), at 19,800 rcf for 15 min. The filter was then washed three times with 150 µl of Buffer UA (15 min at 19,800 rcf). The proteins were reduced with 100 µL of dithiothreitol (DTT, 0.008 M in UA), incubated on a thermomixer at 24 °C for 15 min, and centrifuged 10 min at 19,800 rcf. Filter was further washed with 100 µl of Buffer UA (15 min at 19,800 rcf). Then, proteins were alkylated with 100 µL of iodoacetamide (IAA, 0.05 M iodoacetamide in UA), incubated 20 min at 24 °C in the dark before centrifugation 10 min at 19,800 rcf. Filter was washed with 100 µl of Buffer UA (15 min at 19,800 rcf). The IAA was quenched by the addition of 100 µl of DTT. After an incubation of 15 min at 24 °C, the column was centrifuged 10 min at 19,800 rcf, and further rinsed with 100 µl of Buffer UA. The filter was washed three times with 100 µl of Buffer ABC (0.05 M NH_4_HCO_3_ in water) and centrifuged 10 min at 19,800 rcf. Finally, proteins were digested by trypsin (1 µg, Promega) overnight at 24°C. The digested proteins were recovered by placing the Microcon column on a protein LoBind tube of 1.5 ml followed by centrifugation 10 min at 19,800 rcf. The filter was rinsed with 40 µl of Buffer ABC and centrifuged 10 min at 19,800 rcf. The filtrate was then acidified with a solution of 10% Trifluoroacetic acid (TFA) to reach 0.2% TFA final concentration. The sample was finally dried in a speed-vac up to 20 µl before mass spectrometry analysis (nano-LC-ESI-MS/MS tims TOF Pro, Bruker) coupled with a nanoUHPLC (nanoElute, Bruker).

Peptides were separated by nanoUHPLC on a 75 µm ID, 25 cm C18 column with integrated CaptiveSpray insert (Aurora, Ionopticks, Melbourne) at a flow rate of 200 nL/min, at 50°C. LC mobile phases A was water with 0.1% formic acid (v/v) and B was ACN with formic acid 0.1% (v/v). Samples were loaded directly on the analytical column at a constant pressure of 800 bar. The digest (1 µl) was injected, and the organic content of the mobile phase was increased linearly from 2% B to 15 % in 22 min, from 15 % B to 35% in 38 min, from 35% B to 85% in 3 min. Data acquisition on the tims TOF Pro was performed using Hystar 6.1 and timsControl 2.0. tims TOF Pro data were acquired using 160 ms TIMS accumulation time, mobility (1/K0) range from 0.75 to 1.42 Vs/cm². Mass-spectrometric analysis were carried out using the parallel accumulation serial fragmentation (PASEF) acquisition method (<u>Meier *et al*, 2018</u>). One MS spectra followed by six PASEF MSMS spectra per total cycle of 1.16 s.

All MS/MS samples were analyzed using Mascot (Matrix Science, London, UK; version 2.8.1). Mascot was set up to search the *Brucella abortus* 2308 proteome (3027 entries) downloaded from UniProt (July 2023) and Contaminants_20190304 database assuming the digestion enzyme trypsin. Mascot was searched with a fragment ion mass tolerance of 0.050 Da and a parent ion tolerance of 15 PPM. Carbamidomethyl of cysteine was specified in Mascot as a fixed modification. Deamidated of asparagine and glutamine, oxidation of methionine and acetyl of the n-terminus were specified in Mascot as variable modifications. Scaffold (5.1.1, Proteome Software Inc., Portland, OR) was used to validate MS/MS based peptide and protein identifications. Peptide identifications were accepted if they could be established at greater than 97.0% probability to achieve an FDR less than 1.0% by the Percolator posterior error probability calculation (<u>Kall *et al*, 2008</u>). Protein identifications were accepted if they could be established at greater than 50.0% probability to achieve an FDR less than 1.0% and contained at least 2 identified peptides. Protein probabilities were assigned by the Protein Prophet algorithm (<u>Nesvizhskii *et al*, 2003</u>). Proteins that contained similar peptides and could not be differentiated based on MS/MS analysis alone were grouped to satisfy the principles of parsimony. Proteins sharing significant peptide evidence were grouped into clusters.

## Acknowledgements

We thank M. Loperena Barber and R. Condez-Alvarez lab for their help in OMVs isolation and TLC assay; E. Carlier, M. Waroquier, C. Desy and F. Tilquin for their logistic support; C. Roomans and G. Lima Mendez for her help in the initial steps of protein structure prediction; and G. C. Demazy and S. Burteau for the MS sample preparation; P. Cherry and K. Poncin for the knowledge regarding pull-down; E. Barbieux for the experience of the Tn-seq assay; and staff at UNamur (https://www.unamur.be/) for financial and logistical support. We thank the URBM researchers, Jean-François Collet (UCLouvain) and Basile Beaud Benyahia (Pasteur Institute, Paris) for stimulating discussions. A.L. was supported by a FRIA (FNRS) PhD fellowship. This publication is supported by the Walloon Region as part of the funding for the FRFS-WELBIO strategic axis (X.1512.24). The work was also supported by PDR grants T.0058.20 and T.0068.24 from FRS-FNRS, as well as Concerted Research Action 17/22-087 and 22/27-128 from the Fédération Wallonie-Bruxelles. This research was carried out in the frame of project PID2023-146797OB-C31 financed by MCIN/AEI/10.1303910.13039/501100011033.

## Author contributions

A.R. characterized the *cls*, *olsA* and *olsB* mutants (DOC, infection and TLC), double counting the intra-extra-bacteria in macrophages and helped with the preparation of pull-down assay. M.D. and P.R. performed the proteomic analysis. M.F.G. and A.G.F. performed the lipidomic analysis. R. C. A. contributed to the OMVs isolation and TLC assay. A.L. performed all the other experiments. A.L. and X.D.B. designed the experiments and wrote the manuscript.

The authors declare that they have no conflict of interest.

## Supplementary data

**Table S1.**
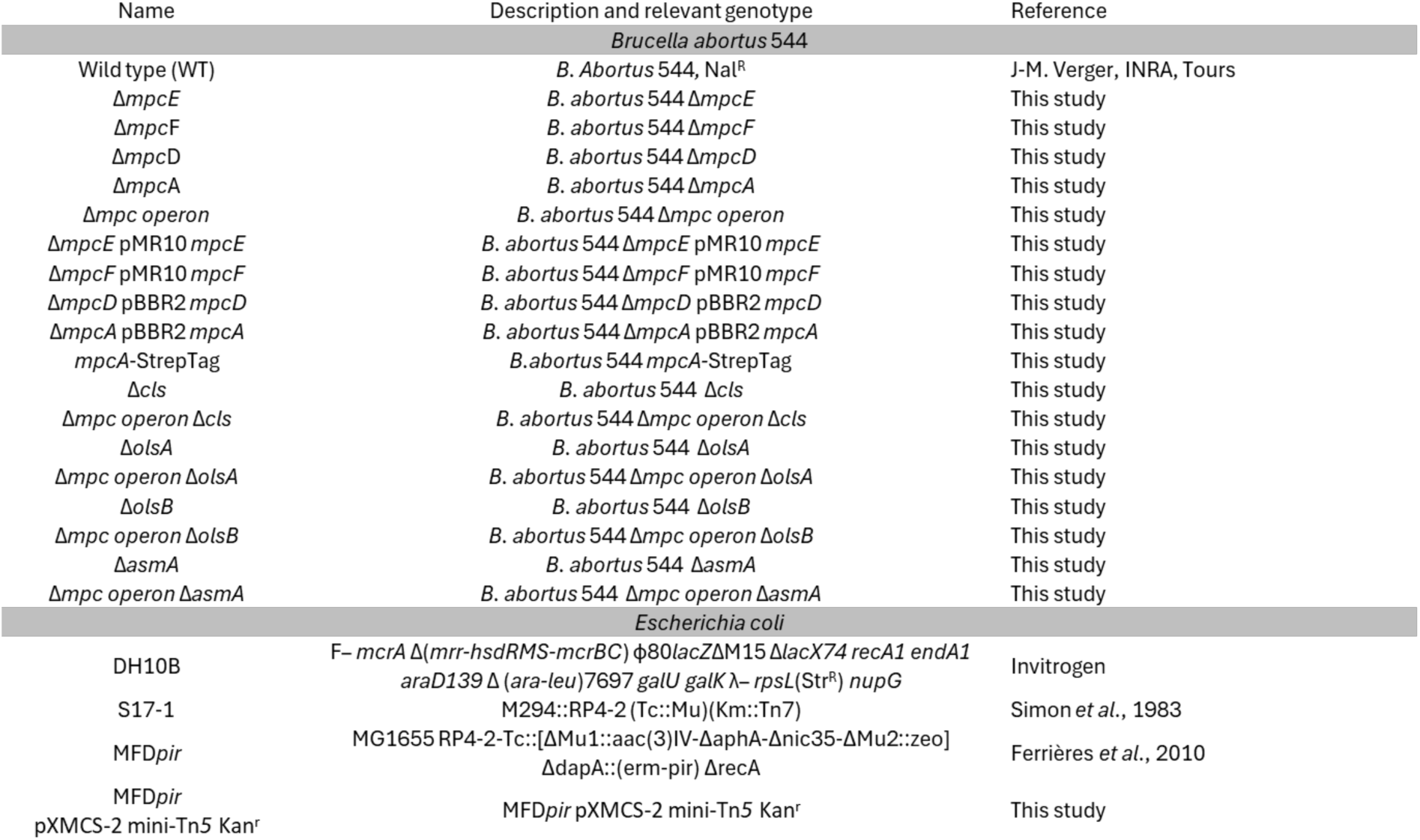
List of strains used in this study.

**Table S2.** List of plasmids used in this study.

| Name | Reference |
| --- | --- |
| pNPTS138 | M. R. K. Alley, Imperial College of Science, London, UK |
| pBBR2 MCS | Deghelt *et al.*, 2014, Unamur, BE |
| pMR10 | C.D. Mohr and R.C. Roberts, Stanford University, US |
| pXMCS-2 mini-Tn5 Kan<sup>r</sup> | Sternon *et al.*, 2018, Unamur, BE |
| pNPTs 138 $\Delta mpcE$ | This study |
| pNPTs 138 $\Delta mpcF$ | This study |
| pNPTs 138 $\Delta mpcD$ | This study |
| pNPTs 138 $\Delta mpcA$ | This study |
| pNPTs 138 $\Delta mpc$ operon | This study |
| pMR10 $\Delta mpcE$ | This study |
| pMR10 $\Delta mpcF$ | This study |
| pBBR2 MCS $\Delta mpcD$ | This study |
| pBBR2 MCS $\Delta mpcA$ | This study |
| pNPTs 138 $mpcA$-Strep Tag | This study |
| pNPTs 138 $\Delta cls$ | This study |
| pJQ200 $\Delta olsA$ | Palacios-Chaves *et al.*, 2011, Universidad de Navarra, Spain |
| pJQ200 $\Delta olsB$ | Palacios-Chaves *et al.*, 2011, Universidad de Navarra, Spain |
| pNPTs 138 $\Delta asmA$ | This study |
| pBBR2 MCS $\Delta asmA$ | G. Potemberg, PhD thesis, Unamur, BE |

**Table S3.**
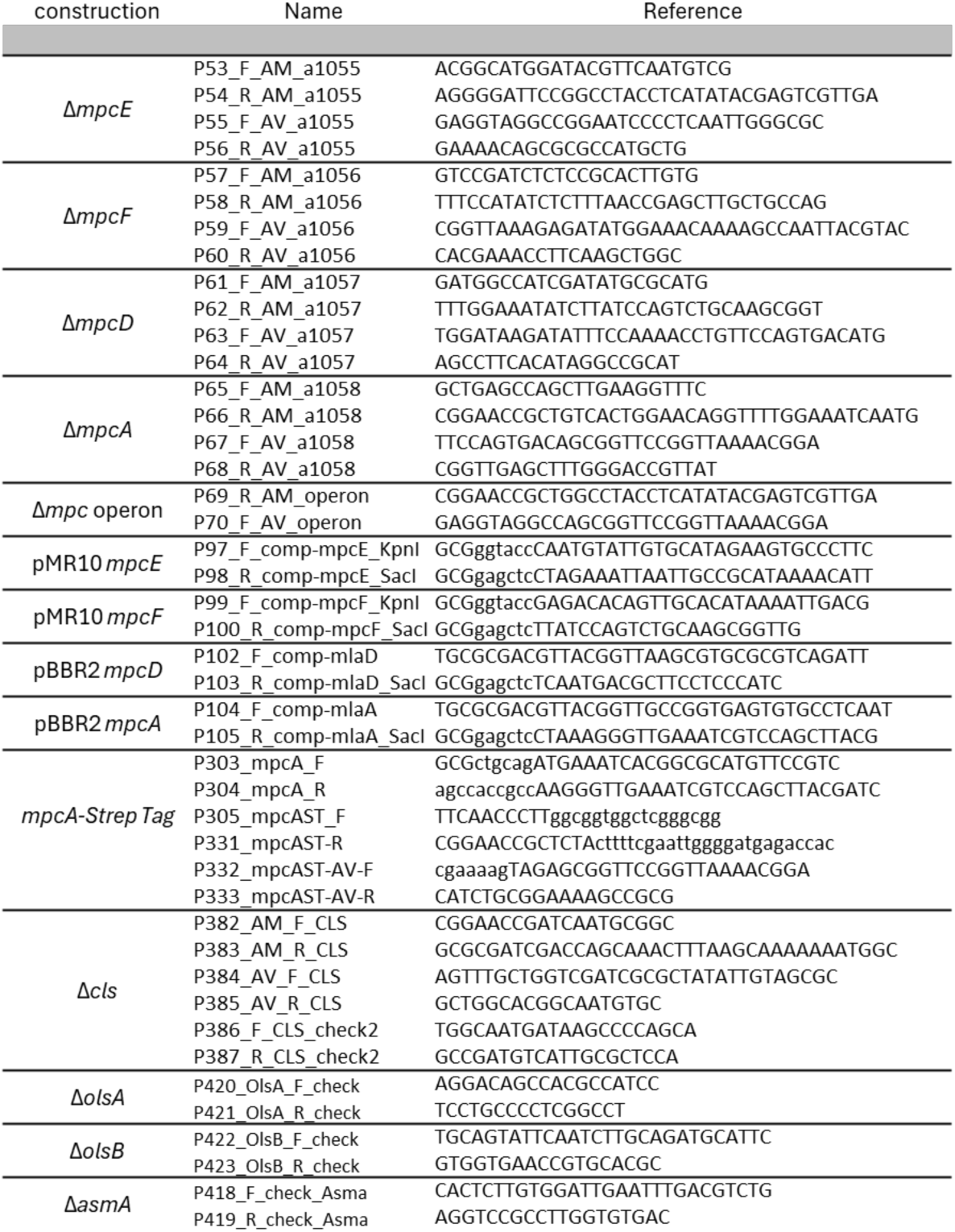
List of primers used in this study.

**Table S4.**
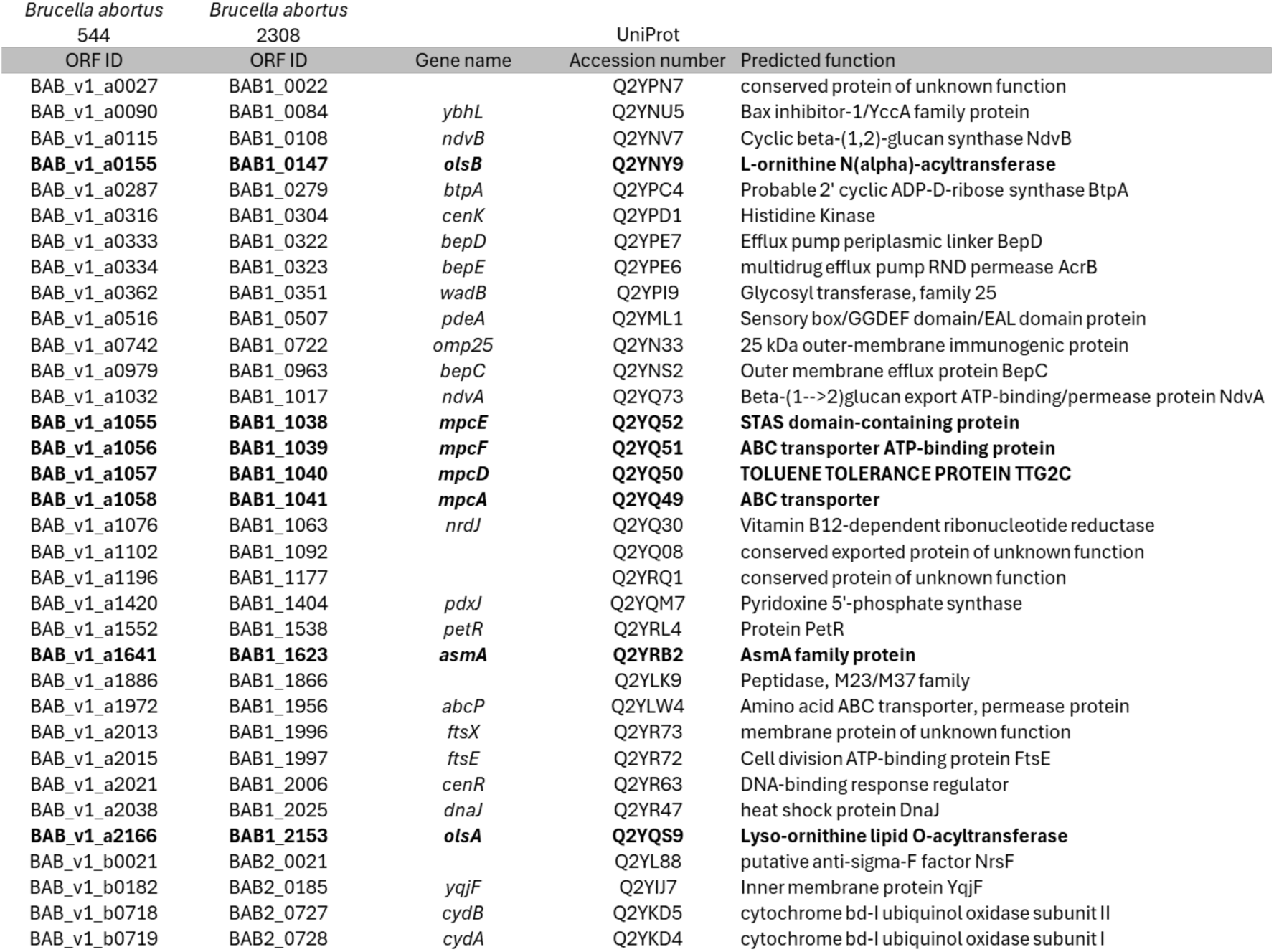
List of ORFs cited in this study.

**Figure S1.**
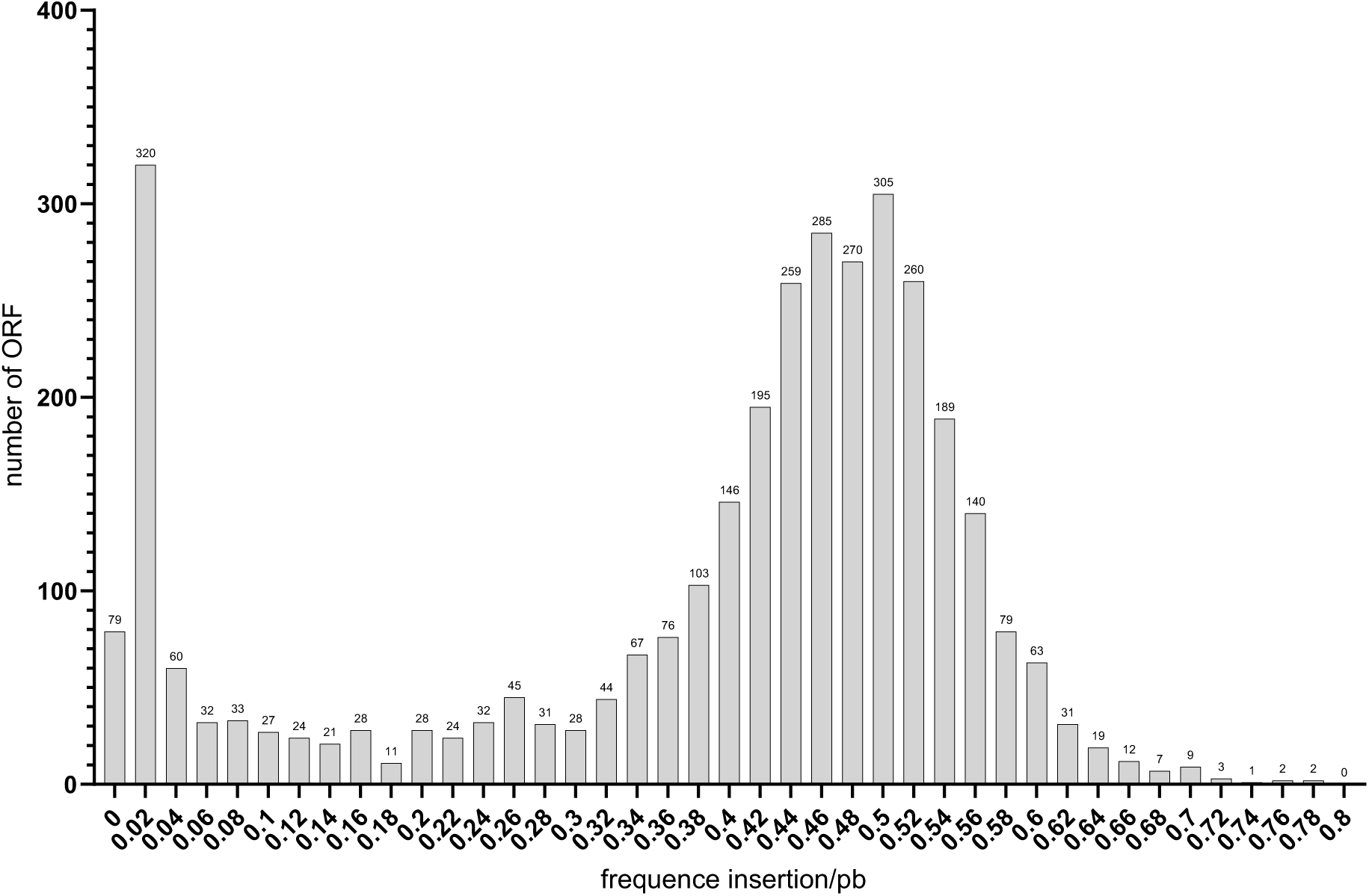
Frequency distribution of the frequence of insertions per pair of bases for mini-Tn*5* across all open reading frames (ORFs) in the *B. abortus* genome. The frequence of insertions per pb for each of 3,390 ORFs from *B. abortus* genome are represented by classes of 0.02. Under 0.1 frequence of insertions per pb, the ORFs are considered essential for growth on TSA rich medium, which correspond to 16.25% of the predicted genes.

**Figure S2.**
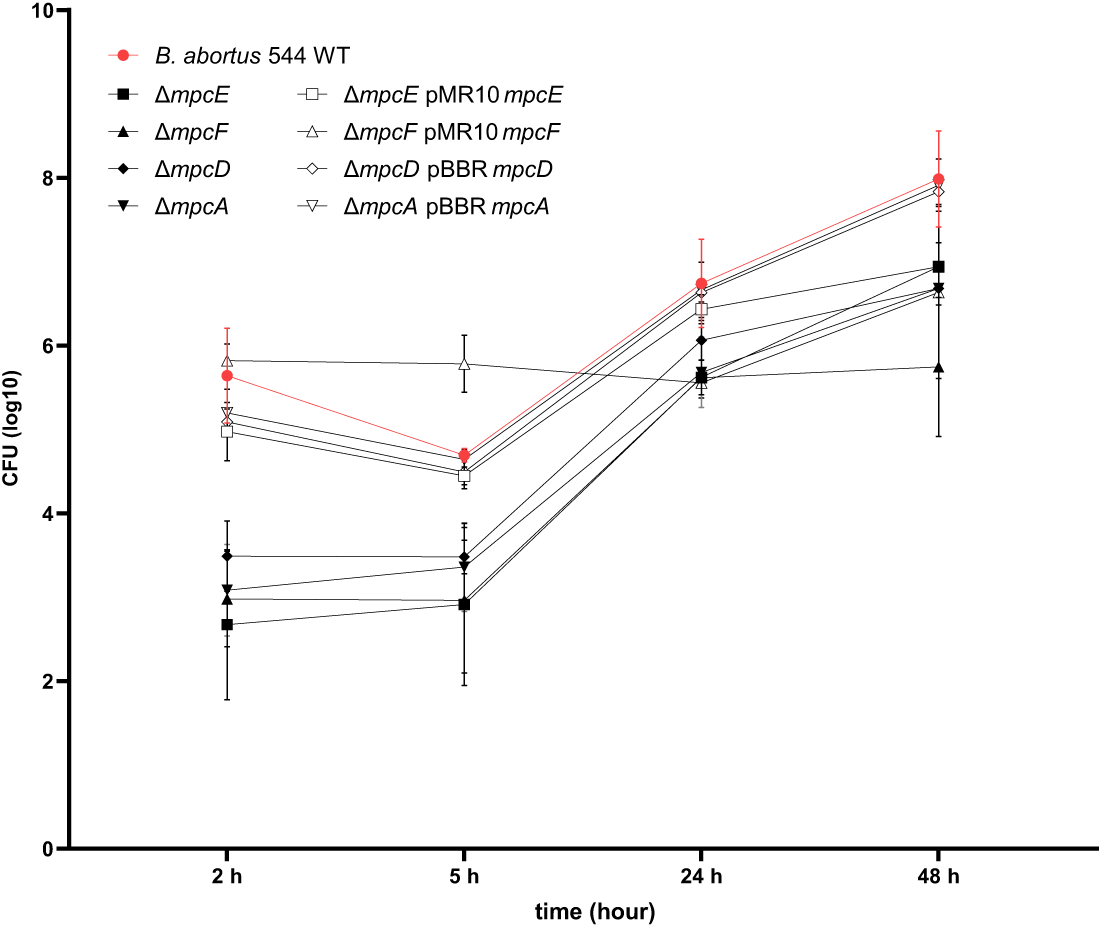
Growth of *mpc* mutants inside the host cell. Intracellular replication of WT, *mpc* mutants and the complemented strains were assessed by counting colony-forming units (CFU) at 2, 5, 24, and 48 hours post-infection of J774.A1 macrophages. The data represents the mean ± SD and were compiled from three independent replicates.

**Figure S3.**
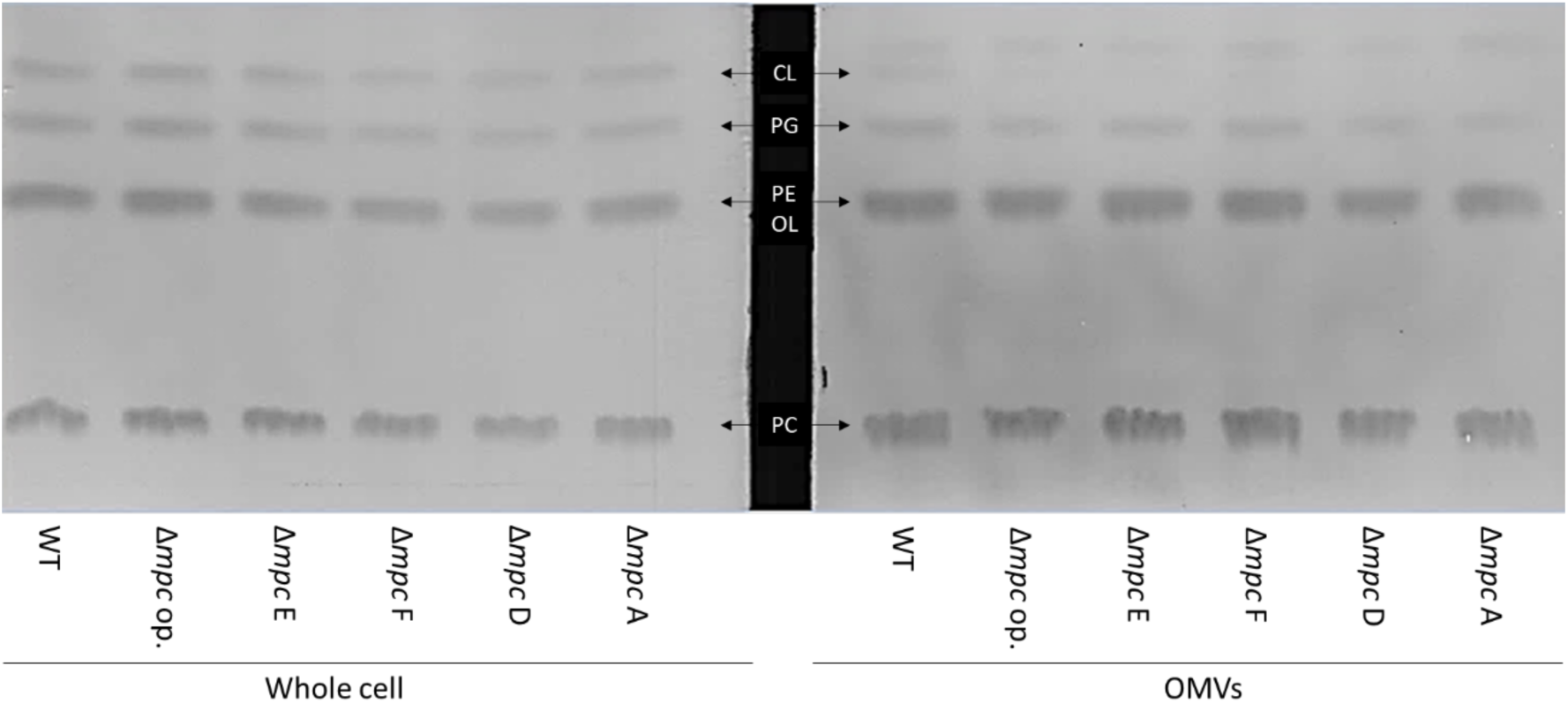
Decreased of cardiolipin (CL) in OMVs in the *mpc* mutants. Thin layer chromatography (TLC) was performed on lipid extracts obtained from both whole cells and outer membrane vesicles (OMVs). The lipids were separated by phosphatidylcholine (PC), phosphatidylethanolamine and ornithine lipids (PE/OL), phosphatidylglycerol (PG) and Cardiolipin (CL) from bottom to the top. In the OMVs samples a band appeared above the CL, but the nature of this lipid is unknown and could not be revealed by staining with iodine vapor.

**Figure S4.**
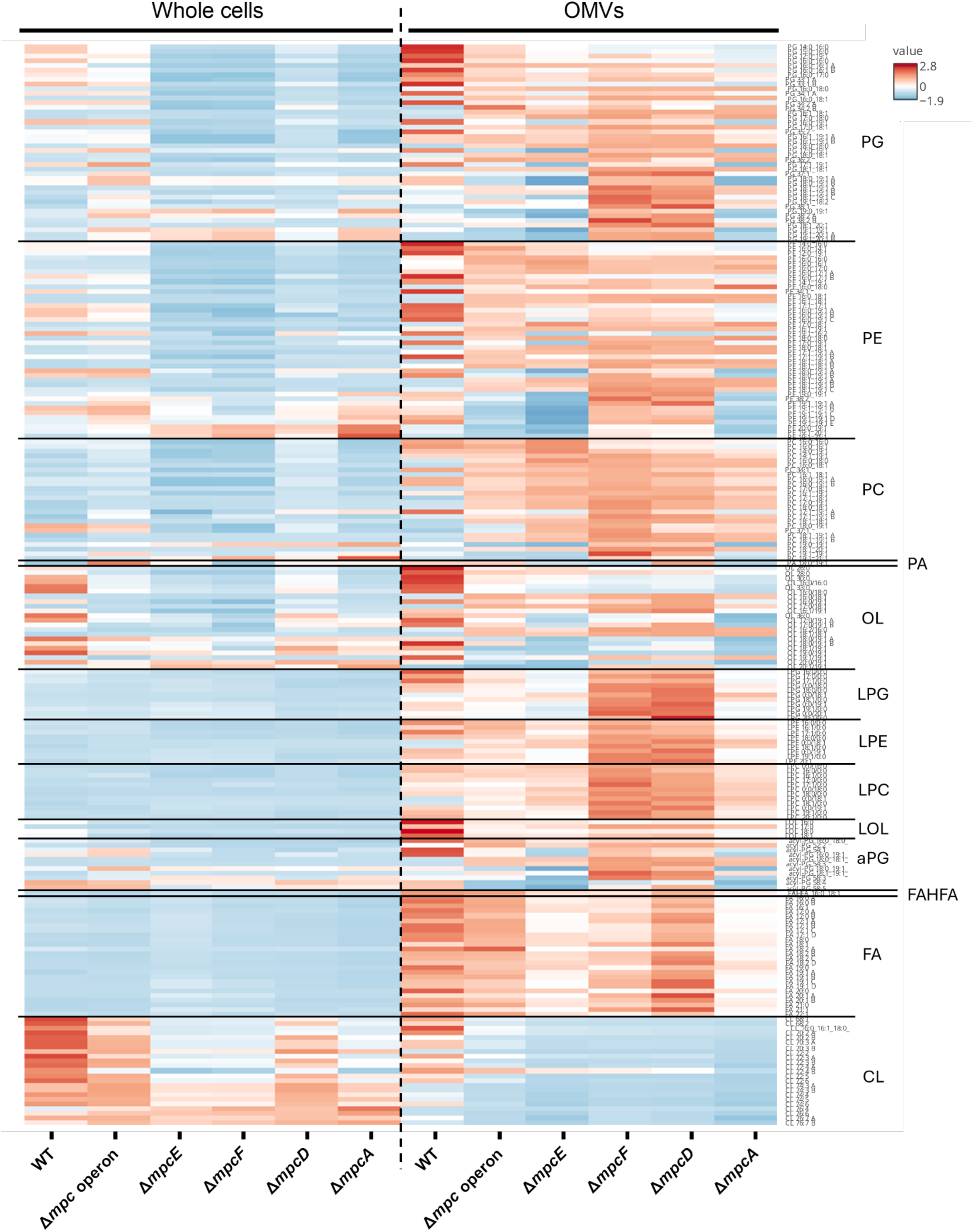
Lipid composition of whole cells compared to outer membrane vesicles. Heat map of abundance of phospholipids identified by mass spectrometry-based lipidomic analysis. PG: phosphatidylglycerol, PE: phosphatidylethanolamine, PC: phosphatidylcholine, PA: Phosphatidate, OL: ornithine lipid, LPG: lyso-phosphatidylglycerol, LPE: lyso-phosphatidylethanolamine, LPC: lyso-phosphatidylcholine, LOL: lyso-ornithine lipid, aPG: aminoacyl-phosphatidylglycerol, FAHFA: fatty acid esters of hydroxy fatty acid, FA: fatty acid, CL: cardiolipin.

**Figure S5.**
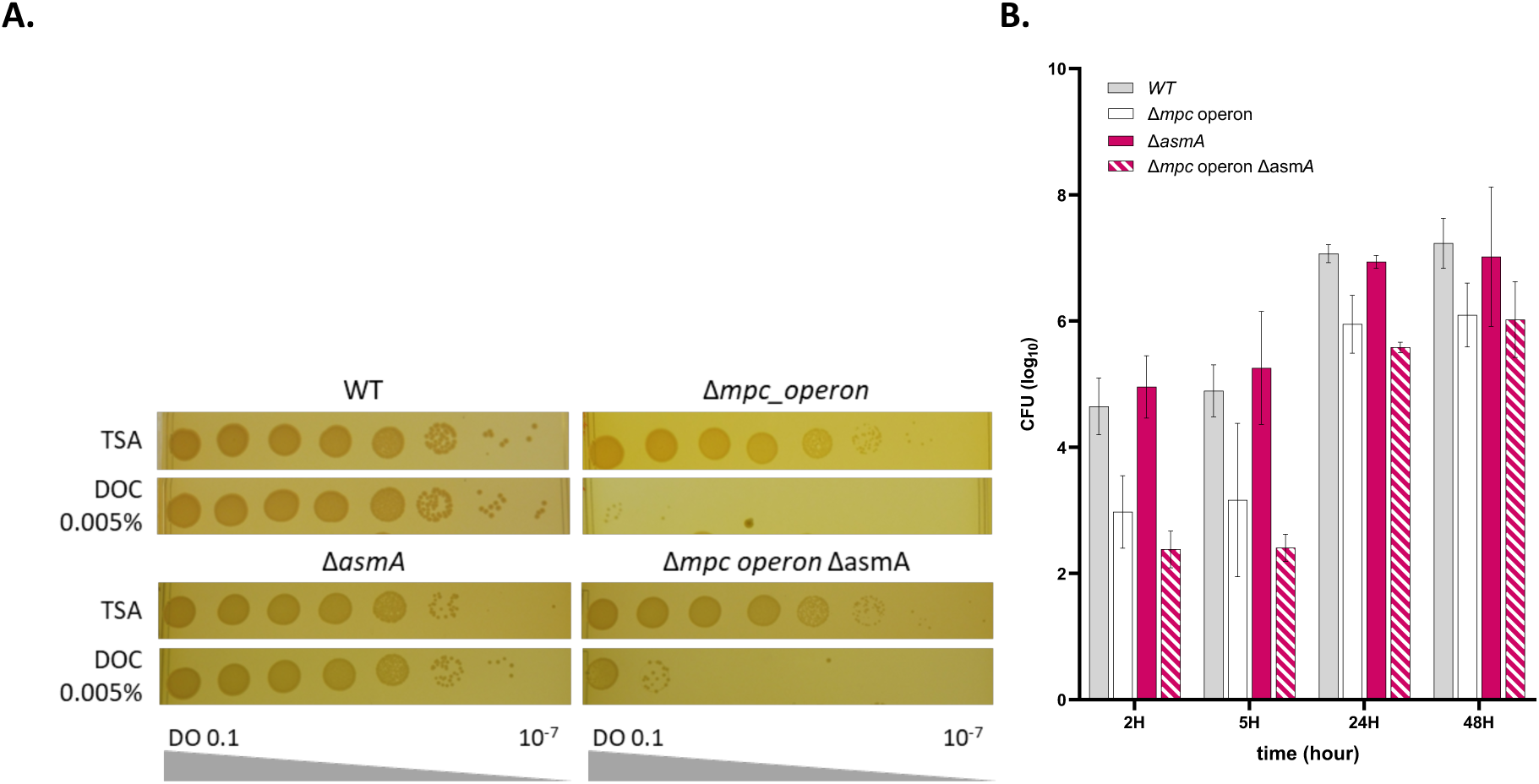
The sensitivity phenotype of *asmA* deletion mutants. **A.** The plating assay on *asmA* mutants, *mpc* operon mutant and double mutants on DOC. All the mutants with deletion of *mpc* operon present the same sensitivity to DOC at 0.005%. The overnight cultures were normalized to 0.1 OD_600_ and serially diluted before being plated onto TSA with or without DOC. **B.** Intracellular replication of WT, *mpc* operon, *asmA* mutant mutant and the double mutants were assessed by CFU at 2, 5, 24, and 48 hours post-infection of J774.A1 macrophages. The data represents the mean ± SD and were compiled from three independent replicates. Statistical significance between the results for a given strain and those for the WT was determined using a two-way ANOVA (not significant: p≥0.05 (not shown); *0.05<p<0.01, **0.01<p<0.001, ***0.001<p<0.0001, ****p< 0.0001).

**Figure S6.**
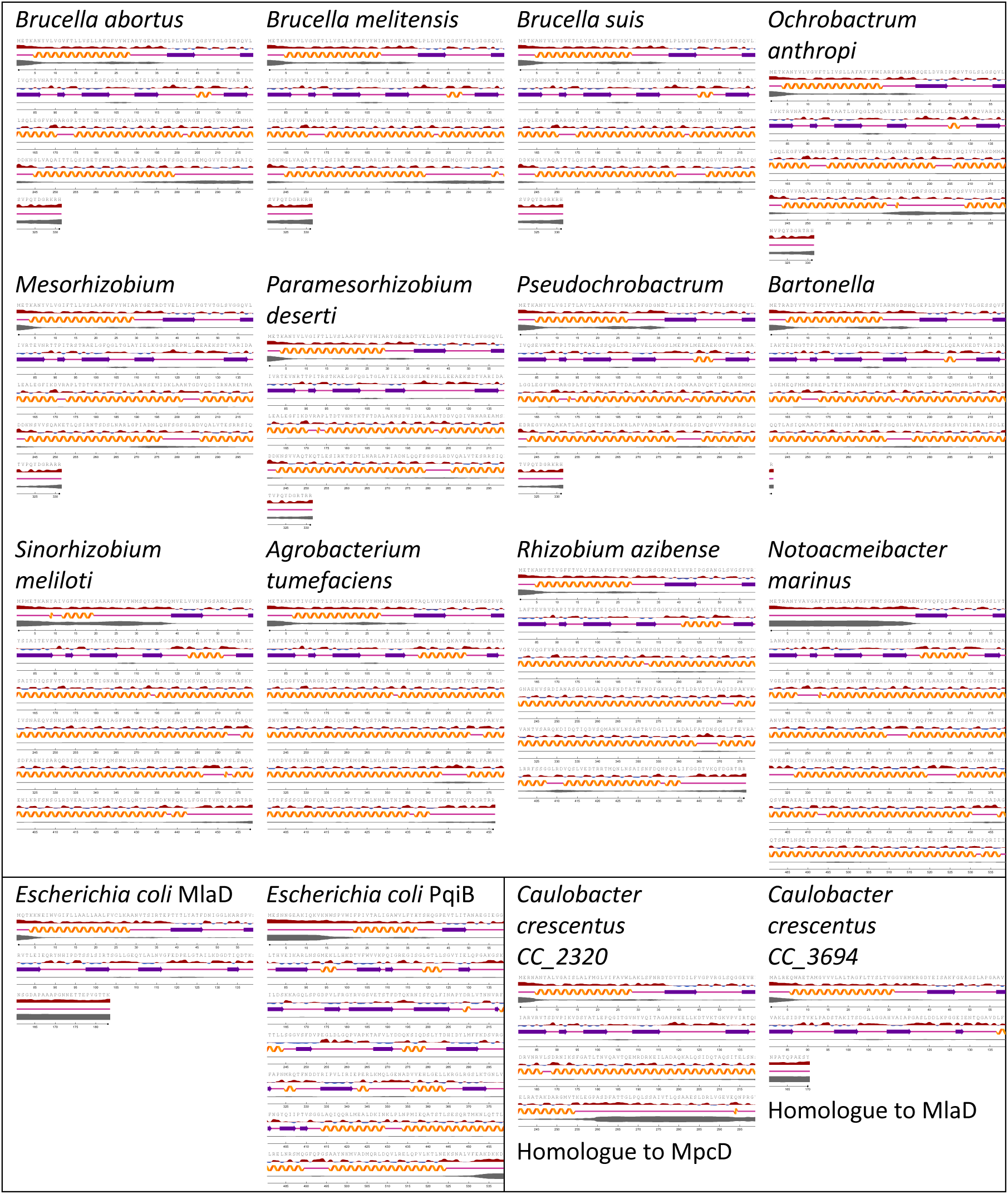
Secondary structure prediction by NetSurfP-2.0 (<u>Klausen *et al*., 2019</u>) of MpcD and homologous proteins in some Hyphomicrobiales bacteria (top panel). Secondary structure of MlaD and PqiB in *E. coli* (lower left panel) and secondary structure prediction of proteins in *Caulobacter crescentus* (lower right panel), one homologous to *B. abortus* MpcD and the other homologous to *E. coli* MlaD.

**Figure S7.**
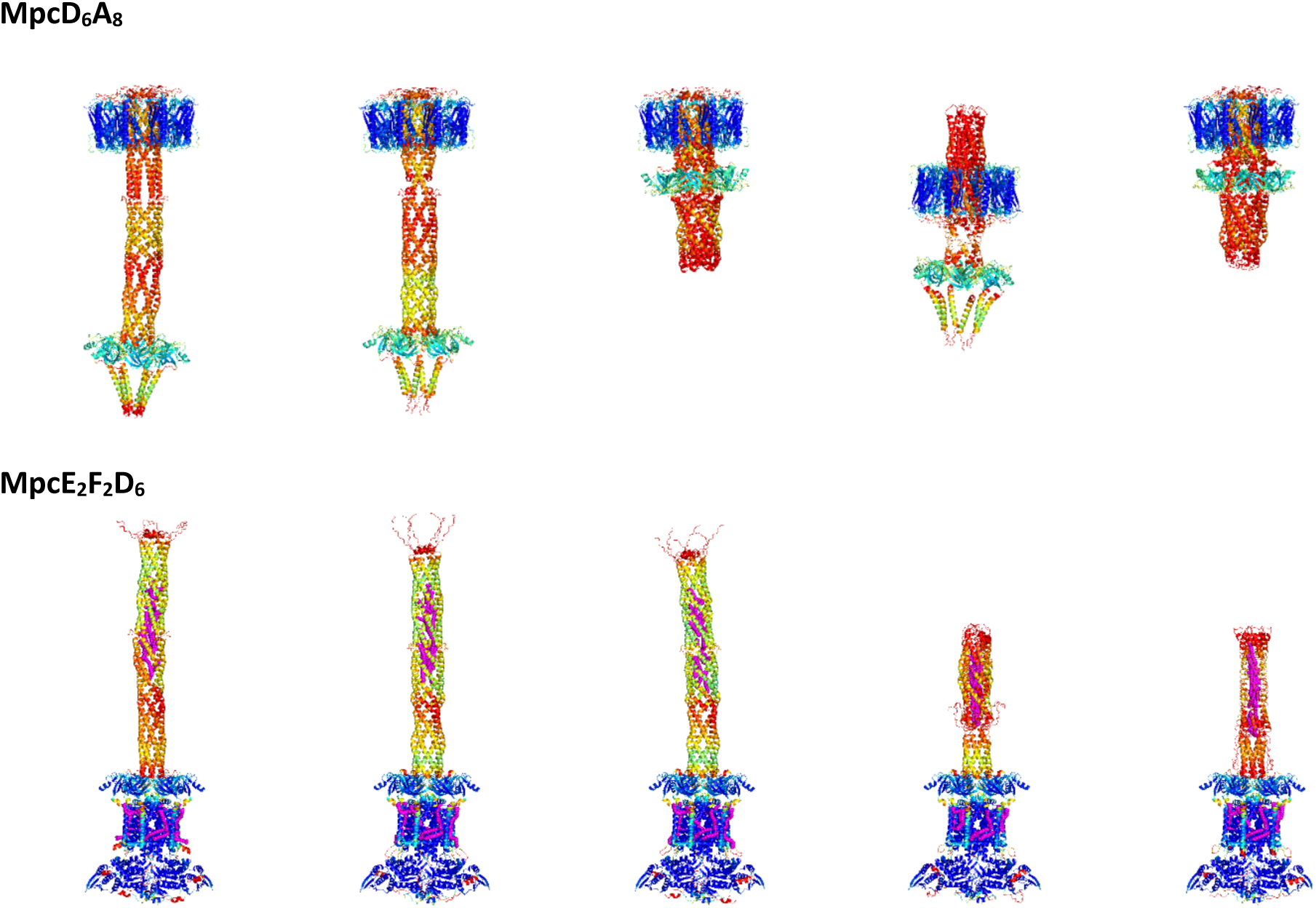
The five models of interaction predicted for the Mpc complex. The AlphaFold3 multimer predictions of biomolecular interactions of complexes of MpcD_6_A_8_ (top) and MpcE_2_F_2_D_6_ (bottom) with 50 oleic acids (in pink) that could potentially mimic the PLs interactions. Coloured with the pLDDT confidence prediction (red pLDDT<50; yellow pLDDT>70; cyan 90>pLDDT>70; blue pLDDT>90). For both predictions, the second model is utilised for the figures.

**Figure S8.**
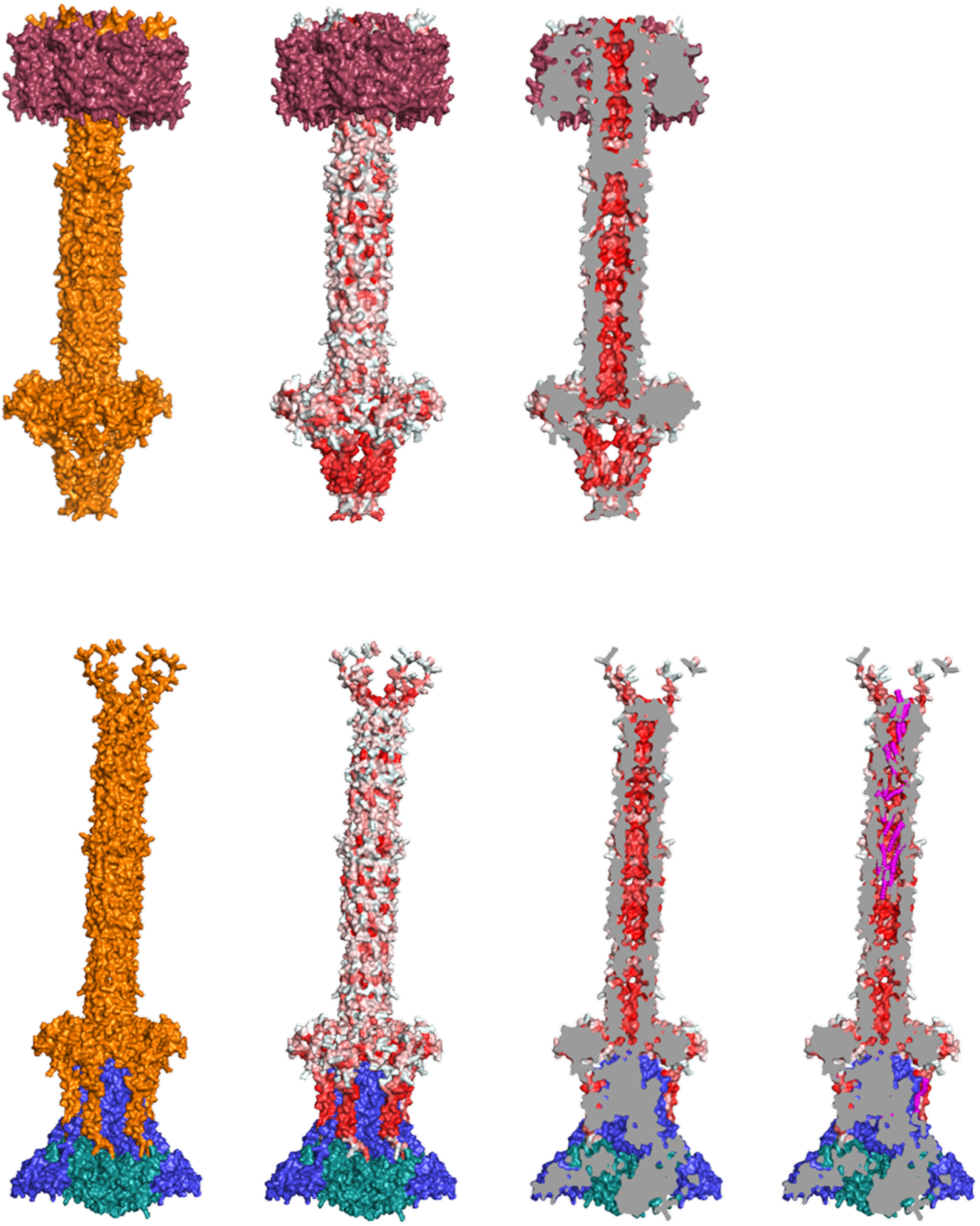
The model predictions for MpcD_6_A_8_ (top) and MpcE_2_F_2_D_6_ (bottom). The surface representation predicted by AlphaFold3 multimer of the biomolecular interactions of complexes of MpcD_6_A_8_ (top) and MpcE_2_F_2_D_6_ (bottom). The initial panel depicts the interaction with the homodimer of MpcE (blue), the homodimer of MpcF (green), the homo-hexamer of MpcD (orange) and the homo-octamer of MpcA (pink). In the second panel, the MpcD homo-hexamer is coloured according to the Eisenberg hydrophobicity scale using the color_h command in PyMol. The third panel presents a slice of the complexes view, allowing observation of the hydrophobicity of the full-length tunnel within the hexameric MpcD channel. The final panel illustrates the slice of the MpcE_2_F_2_D_6_ complex in predicted interaction with 50 oleic acids (magenta), some of which are situated within the channel formed by the coiled-coil alpha-helices of MpcD.

## References

Abellon-Ruiz J, Kaptan SS, Basle A, Claudi B, Bumann D, Kleinekathofer U, van den Berg B (2017) Structural basis for maintenance of bacterial outer membrane lipid asymmetry. Nat Microbiol 2: 1616–1623

Abramson J, Adler J, Dunger J, Evans R, Green T, Pritzel A, Ronneberger O, Willmore L, Ballard AJ, Bambrick J et al (2024) Accurate structure prediction of biomolecular interactions with AlphaFold 3. Nature 630: 493–500

Alakavuklar MA, Fiebig A, Crosson S (2023) The Brucella Cell Envelope. Annu Rev Microbiol

Ando S, Saito M (1987) Chapter 9: TLC and HPTLC of Phospholipids and Glycolipids in Health and Disease. In: *Chromatography of Lipids in Biomedical Research and Clinical Diagnosis*, pp. 266–310.

Arruda S, Bomfim G, Knights R, Huima-Byron T, Riley LW (1993) Cloning of an M. tuberculosis DNA fragment associated with entry and survival inside cells. Science 261: 1454–1457

Awai K, Xu C, Tamot B, Benning C (2006) A phosphatidic acid-binding protein of the chloroplast inner envelope membrane involved in lipid trafficking. Proceedings of the National Academy of Sciences 103: 10817–10822

Bligh EG, Dyer WJ (1959) A rapid method of total lipid extraction and purification. Canadian journal of biochemistry and physiology 37: 911–917

Boratyn GM, Schäffer AA, Agarwala R, Altschul SF, Lipman DJ, Madden TL (2012) Domain enhanced lookup time accelerated BLAST. Biology Direct 7

Brown PJ, de Pedro MA, Kysela DT, Van der Henst C, Kim J, De Bolle X, Fuqua C, Brun YV (2012) Polar growth in the Alphaproteobacterial order Rhizobiales. Proc Natl Acad Sci U S A 109: 1697–1701

Caro-Hernandez P, Fernandez-Lago L, de Miguel MJ, Martin-Martin AI, Cloeckaert A, Grillo MJ, Vizcaino N (2007) Role of the Omp25/Omp31 family in outer membrane properties and virulence of Brucella ovis. Infect Immun 75: 4050–4061

Celli J, de Chastellier C, Franchini DM, Pizarro-Cerda J, Moreno E, Gorvel JP (2003) Brucella evades macrophage killing via VirB-dependent sustained interactions with the endoplasmic reticulum. J Exp Med 198: 545–556

Chen J, Fruhauf A, Fan C, Ponce J, Ueberheide B, Bhabha G, Ekiert DC (2023a) Structure of an endogenous mycobacterial MCE lipid transporter. Nature 620: 445–452

Chen X, Alakavuklar M, A, Fiebig A, Crosson S (2023b) Cross-regulation in a three-component cell envelope stress signaling system of Brucella. mBio 14: e02387–02323

Chong ZS, Woo WF, Chng SS (2015) Osmoporin OmpC forms a complex with MlaA to maintain outer membrane lipid asymmetry in Escherichia coli. Mol Microbiol 98: 1133–1146

Cloeckaert A, Zygmunt MS, Dubray G, Limet JN (1993) Characterization of O-polysaccharide specific monoclonal antibodies derived from mice infected with the rough Brucella melitensis strain B115. Journal of General Microbiology 139: 1551–1556

Cooper BF, Clark R, Kudhail A, Bhabha G, Ekiert DC, Khalid S, Isom GL (2023) Phospholipid transport to the bacterial outer membrane through an envelope-spanning bridge. bioRxiv

Cooper BF, Ratkeviciute G, Clifton LA, Johnston H, Holyfield R, Hardy DJ, Caulton SG, Chatterton W, Sridhar P, Wotherspoon P et al (2024) An octameric PqiC toroid stabilises the outer-membrane interaction of the PqiABC transport system. EMBO Rep 25: 82–101

Coppine J, Kaczmarczyk A, Petit K, Brochier T, Jenal U, Hallez R (2020) Regulation of Bacterial Cell Cycle Progression by Redundant Phosphatases. J Bacteriol 202

Coudray N, Isom GL, MacRae MR, Saiduddin MN, Bhabha G, Ekiert DC (2020) Structure of bacterial phospholipid transporter MlaFEDB with substrate bound. Elife 9

Daniels L, Hanson RS, Phillips J (1993) Chemical analysis. In: Mathods for General and Molecular Bacteriology, Gerhardt P., Murray R.G.E., Wood W.A., Krieg N.R. (eds.) pp. 512-554. ASM: Washington, D.C.

Deghelt M, Mullier C, Sternon JF, Francis N, Laloux G, Dotreppe D, Van der Henst C, Jacobs-Wagner C, Letesson JJ, De Bolle X (2014) G1-arrested newborn cells are the predominant infectious form of the pathogen Brucella abortus. microbial cell 5: 4366

Ekiert DC, Bhabha G, Isom GL, Greenan G, Ovchinnikov S, Henderson IR, Cox JS, Vale RD (2017) Architectures of Lipid Transport Systems for the Bacterial Outer Membrane. Cell 169: 273–285 e217

Fernández-García M, Ares-Arroyo M, Wedel E, Montero N, Barbas C, Rey-Stolle MF, González-Zorn B, García A (2023) Multiplatform Metabolomics Characterization Reveals Novel Metabolites and Phospholipid Compositional Rules of Haemophilus influenzae Rd KW20. International Journal of Molecular Sciences 24

Gao JL, Weissenmayer B, Taylor AM, Thomas-Oates J, Lopez-Lara IM, Geiger O (2004) Identification of a gene required for the formation of lyso-ornithine lipid, an intermediate in the biosynthesis of ornithine-containing lipids. Mol Microbiol 53: 1757–1770

Giacometti SI, MacRae MR, Dancel-Manning K, Bhabha G, Ekiert DC (2022) Lipid Transport Across Bacterial Membranes. Annu Rev Cell Dev Biol 38: 125–153

Godessart P, Lannoy A, Dieu M, Van der Verren SE, Soumillion P, Collet JF, Remaut H, Renard P, De Bolle X (2021) beta-Barrels covalently link peptidoglycan and the outer membrane in the alpha-proteobacterium Brucella abortus. Nat Microbiol 6: 27–33

Grabowicz M (2018) Lipoprotein Transport: Greasing the Machines of Outer Membrane Biogenesis: Re-Examining Lipoprotein Transport Mechanisms Among Diverse Gram-Negative Bacteria While Exploring New Discoveries and Questions. Bioessays 40: e1700187

Grasekamp KP, Beaud Benyahia B, Taib N, Audrain B, Bardiaux B, Rossez Y, Izadi-Pruneyre N, Lejeune M, Trivelli X, Chouit Z et al (2023) The Mla system of diderm Firmicute Veillonella parvula reveals an ancestral transenvelope bridge for phospholipid trafficking. Nat Commun 14: 7642

Isom GL, Coudray N, MacRae MR, McManus CT, Ekiert DC, Bhabha G (2020) LetB Structure Reveals a Tunnel for Lipid Transport across the Bacterial Envelope. Cell 181: 653–664 e619

Isom GL, Davies NJ, Chong ZS, Bryant JA, Jamshad M, Sharif M, Cunningham AF, Knowles TJ, Chng SS, Cole JA et al (2017) MCE domain proteins: conserved inner membrane lipid-binding proteins required for outer membrane homeostasis. Sci Rep 7: 8608

Kall L, Storey JD, Noble WS (2008) Non-parametric estimation of posterior error probabilities associated with peptides identified by tandem mass spectrometry. Bioinformatics 24: i42–48

Klausen MS, Jespersen MC, Nielsen H, Jensen KK, Jurtz VI, Sonderby CK, Sommer MOA, Winther O, Nielsen M, Petersen B et al (2019) NetSurfP-2.0: Improved prediction of protein structural features by integrated deep learning. Proteins 87: 520–527

Köfeler HC, Eichmann TO, Ahrends R, Bowden JA, Danne-Rasche N, Dennis EA, Fedorova M, Griffiths WJ, Han X, Hartler J et al (2021) Quality control requirements for the correct annotation of lipidomics data. Nature Communications 12

Konovalova A, Kahne DE, Silhavy TJ (2017) Outer Membrane Biogenesis. Annu Rev Microbiol 71: 539–556

Kumar S, Ruiz N (2023) Bacterial AsmA-Like Proteins: Bridging the Gap in Intermembrane Phospholipid Transport. Contact (Thousand Oaks) 6: 25152564231185931

Laine CG, Johnson VE, Scott HM, Arenas-Gamboa AM (2023) Global Estimate of Human Brucellosis Incidence. Emerg Infect Dis 29: 1789–1797

Lakey BD, Myers KS, Alberge F, Mettert EL, Kiley PJ, Noguera DR, Donohue TJ (2022) The essential Rhodobacter sphaeroides CenKR two-component system regulates cell division and envelope biosynthesis. PLoS Genet 18: e1010270

Lestrate P, Delrue RM, Danese I, Didembourg C, Taminiau B, Mertens P, De Bolle X, Tibor A, Tang CM, Letesson JJ (2000) Identification and characterization of in vivo attenuated mutants of Brucella melitensis. Mol Microbiol 38: 543–551

Liu W, Dong H, Liu W, Gao X, Zhang C, Wu Q (2012) OtpR regulated the growth, cell morphology of B. melitensis and tolerance to beta-lactam agents. Vet Microbiol 159: 90–98

Lu B, Benning C (2009) A 25-amino acid sequence of the Arabidopsis TGD2 protein is sufficient for specific binding of phosphatidic acid. J Biol Chem 284: 17420–17427

Malinverni JC, Silhavy TJ (2009) An ABC transport system that maintains lipid asymmetry in the gram-negative outer membrane. Proc Natl Acad Sci U S A 106: 8009–8014

Martin FA, Posadas DM, Carrica MC, Cravero SL, O’Callaghan D, Zorreguieta A (2009) Interplay between two RND systems mediating antimicrobial resistance in Brucella suis. J Bacteriol 191: 2530–2540

Martínez de Tejada G, Pizarro-Cerdá J, Moreno E, Moriyón I (1995) The outer membranes of Brucella spp. are resistant to bactericidal cationic peptides. Infection and immunity 63: 3054–3061

Meier F, Brunner AD, Koch S, Koch H, Lubeck M, Krause M, Goedecke N, Decker J, Kosinski T, Park MA et al (2018) Online Parallel Accumulation-Serial Fragmentation (PASEF) with a Novel Trapped Ion Mobility Mass Spectrometer. Mol Cell Proteomics 17: 2534–2545

Moreno E, Moriyón I (2006) The Genus Brucella. In: *The Prokaryotes*, pp. 315–456.

Moriyon I, Berman DT (1982) Effects of nonionic, ionic, and dipolar ionic detergents and EDTA on the Brucella cell envelope. Journal of Bacteriology 152: 822–828

Nakayama T, Zhang-Akiyama QM (2017) pqiABC and yebST, Putative mce Operons of Escherichia coli, Encode Transport Pathways and Contribute to Membrane Integrity. J Bacteriol 199

Nesvizhskii AI, Keller A, Kolker E, Aebersold R (2003) A Statistical Model for Identifying Proteins by Tandem Mass Spectrometry. Analytical Chemistry 75: 4646–4658

Okuda S, Sherman DJ, Silhavy TJ, Ruiz N, Kahne D (2016) Lipopolysaccharide transport and assembly at the outer membrane: the PEZ model. Nature Reviews Microbiology 14: 337–345

Palacios-Chaves L, Conde-Alvarez R, Gil-Ramirez Y, Zuniga-Ripa A, Barquero-Calvo E, Chacon-Diaz C, Chaves-Olarte E, Arce-Gorvel V, Gorvel JP, Moreno E et al (2011) Brucella abortus ornithine lipids are dispensable outer membrane components devoid of a marked pathogen-associated molecular pattern. PLoS One 6: e16030

Porte F, Liautard J-P, Köhler S (1999) Early acidification of phagosomes containing Brucella suis is essential for intracellular survival in murine macrophages. infect Immun 67: 4041–4047

Posadas DM, Martin FA, Sabio y Garcia JV, Spera JM, Delpino MV, Baldi P, Campos E, Cravero SL, Zorreguieta A (2007) The TolC homologue of Brucella suis is involved in resistance to antimicrobial compounds and virulence. Infect Immun 75: 379–389

Potemberg G, Demars A, Barbieux E, Reboul A, Stubbe FX, Galia M, Lagneaux M, Comein A, Denis O, Perez-Morga D et al (2022) Genome-wide analysis of Brucella melitensis genes required throughout intranasal infection in mice. PLoS Pathog 18: e1010621

Sawyer J, Martin N, Hancock R (1988) Interaction of macrophage cationic proteins with the outer membrane of Pseudomonas aeruginosa. infect Immun 55: 693–698

Schindelin J, Arganda-Carreras I, Frise E, Kaynig V, Longair M, Pietzsch T, Preibisch S, Rueden C, Saalfeld S, Schmid B et al (2012) Fiji: an open-source platform for biological-image analysis. Nat Methods 9: 676–682

Servais C, Vassen V, Verhaeghe A, Kuster N, Carlier E, Phegnon L, Mayard A, Auberger N, Vincent S, De Bolle X (2023) Lipopolysaccharide biosynthesis and traffic in the envelope of the pathogen Brucella abortus. Nat Commun 14: 911

Shrivastava R, Chng SS (2019) Lipid trafficking across the Gram-negative cell envelope. J Biol Chem 294: 14175–14184

Silhavy TJ, Kahne D, Walker S (2010) The bacterial cell envelope. Cold Spring Harb Perspect Biol 2: a000414

Skerker JM, Prasol MS, Perchuk BS, Biondi EG, Laub MT (2005) Two-component signal transduction pathways regulating growth and cell cycle progression in a bacterium: a system-level analysis. PLoS Biol 3: e334

Starr T, Ng TW, Wehrly TD, Knodler LA, Celli J (2008) Brucella intracellular replication requires trafficking through the late endosomal/lysosomal compartment. Traffic 9: 678–694

Sternon JF, Godessart P, Goncalves de Freitas R, Van der Henst M, Poncin K, Francis N, Willemart K, Christen M, Christen B, Letesson JJ et al (2018) Transposon Sequencing of Brucella abortus Uncovers Essential Genes for Growth In Vitro and Inside Macrophages. Infect Immun 86

Tang X, Chang S, Qiao W, Luo Q, Chen Y, Jia Z, Coleman J, Zhang K, Wang T, Zhang Z et al (2021) Structural insights into outer membrane asymmetry maintenance in Gram-negative bacteria by MlaFEDB. Nat Struct Mol Biol 28: 81–91

Tomasek D, Kahne D (2021) The assembly of beta-barrel outer membrane proteins. Curr Opin Microbiol 60: 16–23

Urdaneta V, Casadesus J (2017) Interactions between Bacteria and Bile Salts in the Gastrointestinal and Hepatobiliary Tracts. Front Med (Lausanne) 4: 163

Vassen V, Valotteau C, Feuillie C, Formosa-Dague C, Dufrene YF, De Bolle X (2019) Localized incorporation of outer membrane components in the pathogen Brucella abortus. EMBO J 38

Wotherspoon P, Johnston H, Hardy DJ, Holyfield R, Bui S, Ratkeviciute G, Sridhar P, Colburn J, Wilson CB, Colyer A et al (2024) Structure of the MlaC-MlaD complex reveals molecular basis of periplasmic phospholipid transport. Nat Commun 15: 6394

Zhang X, Ferguson-Miller SM, Reid GE (2011) Characterization of ornithine and glutamine lipids extracted from cell membranes of Rhodobacter sphaeroides. Journal of the American Society for Mass Spectrometry 20: 198–212

Zhang X, Ren J, Li N, Liu W, Wu Q (2009) Disruption of the BMEI0066 gene attenuates the virulence of Brucella melitensis and decreases its stress tolerance. International Journal of Biological Sciences 5: 570–577

Zheng L, Lin Y, Lu S, Zhang J, Bogdanov M (2017) Biogenesis, transport and remodeling of lysophospholipids in Gram-negative bacteria. Biochim Biophys Acta Mol Cell Biol Lipids 1862: 1404–1413

